# Evaluation role of miR-124 in neurodegenerative diseases: literature review and in silico analysis

**DOI:** 10.1101/2021.10.17.464692

**Authors:** Javad Amini, Bahram Bibak, Amir R Afshar, Amirhossein Sahebkar

## Abstract

Neurodegenerative diseases (ND) are characterized by loss of function and structure of neurons. NDs like Alzheimer’s disease (AD) and Parkinson’s disease (PD) have high burden on the society and patients. Currently microRNAs (miRNAs) approach is growing. miRNAs express in different tissues, especially in the central neuron systems (CNS). miRNAs have a dynamic role in the CNS among this miRNAs, miR-124 significantly express in the CNS. Studies on miR-124 have shown that miR-124 improves ND. In this study, we evaluated the role of miR-124 in the ND by literature review and in silico analysis. We used Pubmed database to find miR-124 function in the Alzheimer’s disease, Parkinson’s disease, Multiple sclerosis, Huntington’s disease and amyotrophic lateral sclerosis. To better understand the role of miR-124 in the neurons, RNA-seq data form miR-124-deleted neuronal cells extracted from GEO database and analyzed in Galaxy platform. According literature review miR-124 attenuates inflammation and apoptosis in the ND by target NF-kb signaling pathway and regulation of BAX/BCL-2. miR-124 targets BACE1 and decreases level of Aβ. RNA-seq data showed miR-124 downregulation, an increase in chemokine gene like CCL1 and cytokine-cytokine receptor-interaction, as well as MAPK-signaling pathway. Our study shows that miR-124 can be promising therapeutic approaches to ND.

## Introduction

Neurodegenerative diseases (ND) are characterized by chronic progressive loss of the structure and functions of neuronal materials in the central nervous system (1). ND is a major threat for human health and its rate escalates the elderly population increases. ND include Alzheimer’s disease (AD), Parkinson’s disease (PD), Huntington’s disease (HD), amyotrophic lateral sclerosis (ALS), frontotemporal dementia (FTD), and the spinocerebellar ataxias are all different in their pathophysiology. Overall, it causes memory and cognitive impairments and it also affects a person’s ability to move, speak and breathe (2). AD is the most common cause of dementia and is characterized by early and progressive memory deficit as well as cognitive disorder (3). The worldwide prevalence of just AD in 2015 is estimated at 44 million and it is predicted to double by 2050 (4). Several common risk factors of ND are exposure to environmental pollutants, oxidative stress, aging, head trauma, and protein dysfunction (5). Recently, coast burden of AD in the USA has reached nearly $250 billion (USD) annually and it is estimated to rise to $1 at 2050 while more than 99% of all type of AD therapies failed in disease-modifying therapies (6). PD is the most common movement disorder and the second most common ND. 1-2 per 1000 population have PD and it increases to 1% for the population above 60 years (7). In AD, PD has both genetic and sporadic types, but in FTD, genetic factors are stronger, and almost all cases of HD are inherited (8).

In recent years, microRNA (miRNA) approach has been growing fast. miRNAs are implicated in the pathogenesis of several diseases such as ND and they were introduced as candidates for diagnostic and prognostic biomarkers, both as predictors of drug response and as therapeutic agents (9). miRNAs are found in bodily fluids such as blood, cerebrospinal fluid (CSF) and tears. miRNAs have an important role in CNS. They can act as a messenger, as well as developmental and differentiation regulator (10).

miR-124 is one of high express miRNA in the brain. miRNA-124 is involved in neurogenesis, synapse morphology, neurotransmission, inflammation, autophagy and mitochondrial function in the brain (11). Interestingly, miR-124 are almost not expressed in non-nervous tissues. miR-124 expression is in different parts of the brain, such as cortex, cerebellum and with mater (12) (Figure 1). Abnormal expression of miR-124 discovered in neural disease includes AD, PD, HD and hypoxic-Ischemic Encephalopathy and ischemic stroke and expression of miR-124 improves ND (13). Moreover, miR-124 protects traumatically spinal cord injury via inhibition of M1 microglia and A1 astrocytes (14).

**Figure 1.**
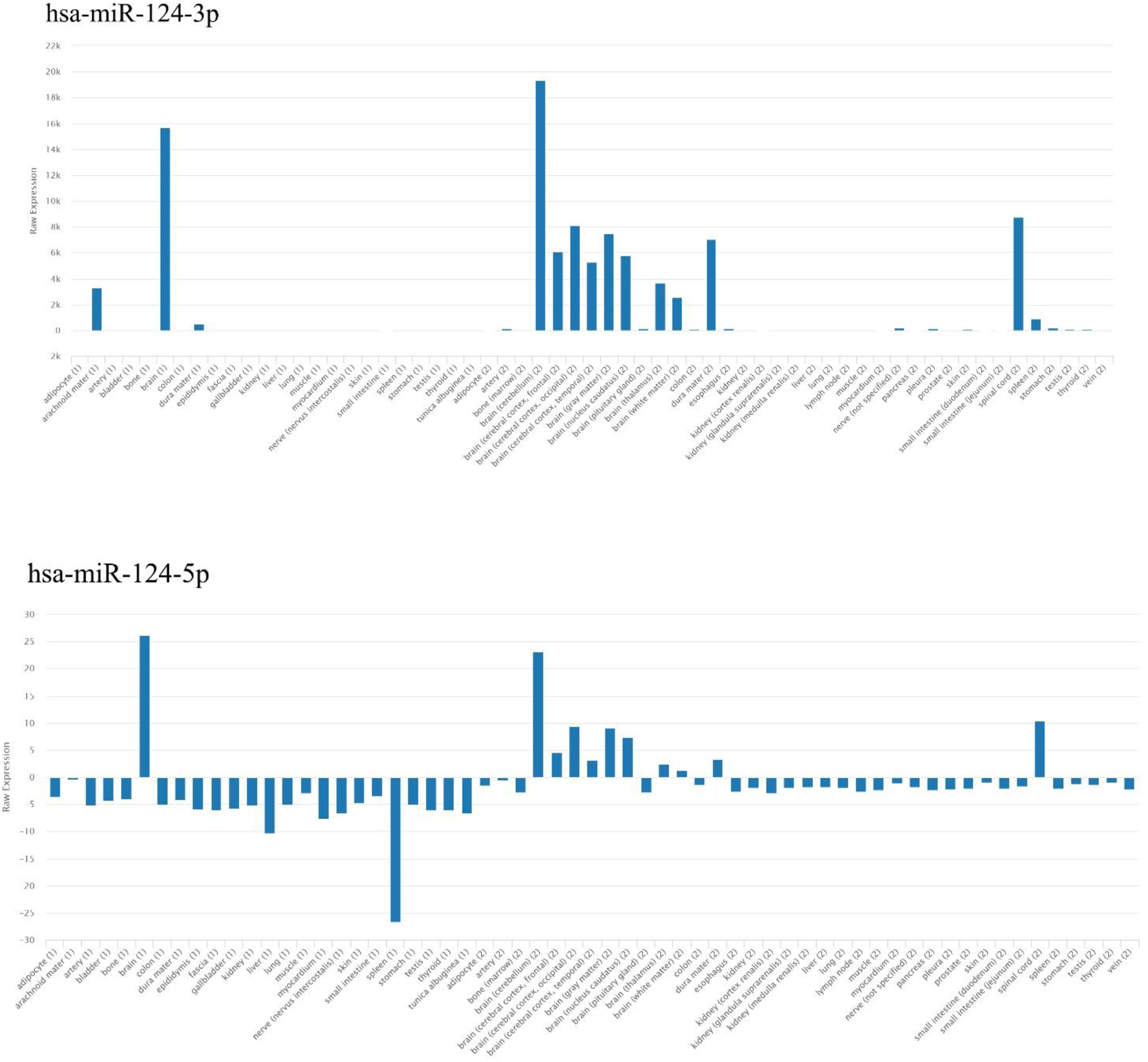
Overview of miR-124 expression in the different tissue in the human.

In this study, we evaluated miR-124’s role in the neurodegenerative disease by literature review and in silico analysis. First of all, we literature reviewed studies on miR-124 in ND. Then, RNA-seq data employed for find role of miR-124 in neuronal cells.

## Material and methods

In this study, we used “miR-124 Alzheimer’s disease”, “miR-124 Parkinson’s disease”, “miR-124 Huntington disease”, “miR-124 multiple sclerosis” and “mir-124 amyotrophic lateral sclerosis” as the keyword in Pubmed database to 2021 may 31 to obtain miR-124’s role in ND. After removing duplication and scanning articles, we selected miRNA studies on brain tissue. According to preferred reporting items for systematic reviews and meta-analyses (PRISMA) instruction, we merged the remaining titles across search databases (Figure 2).

**Figure 2.**
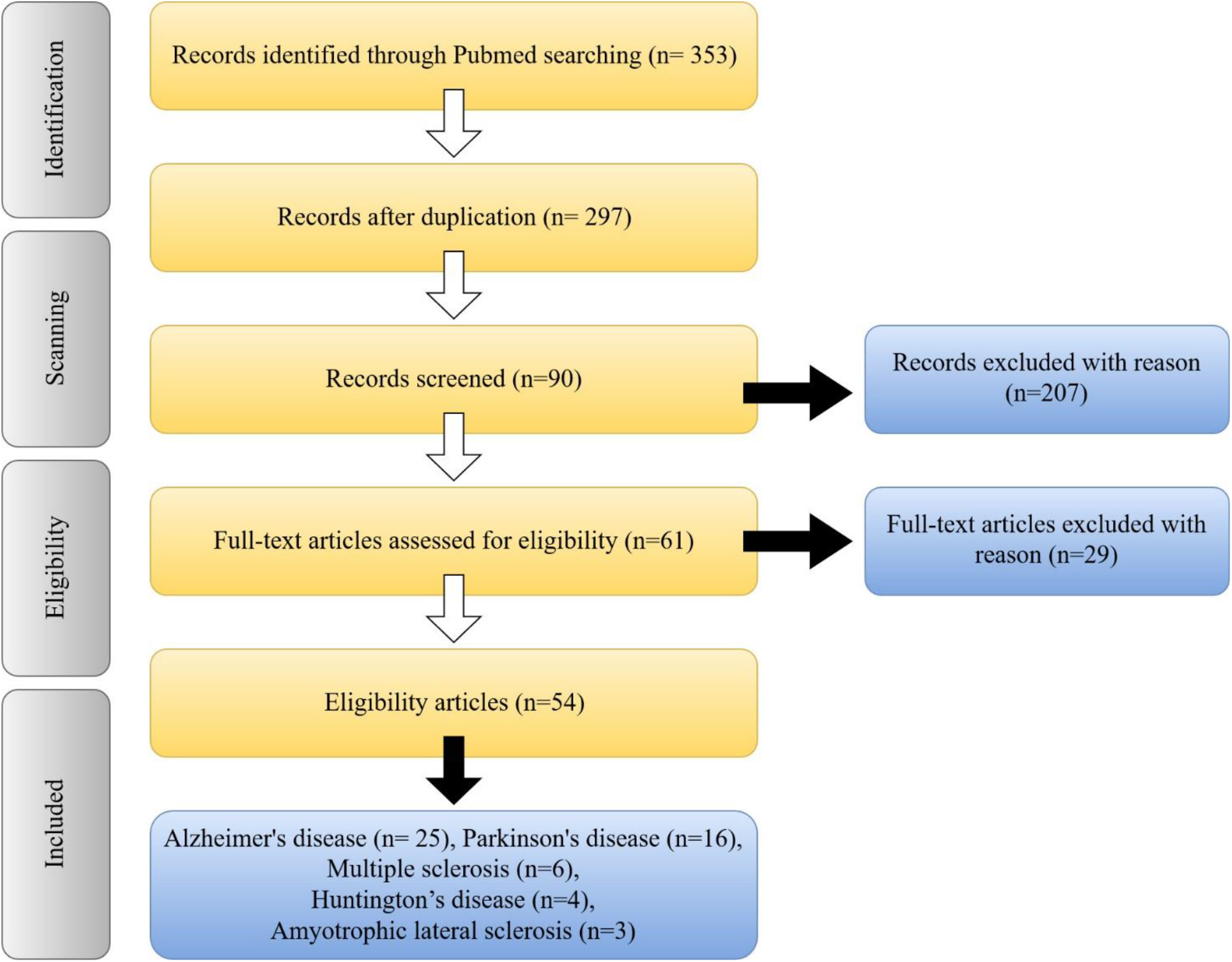
Overview of search methodology.

### RNA-seq data

The Gene Expression Omnibus (GEO) is a public database for gene expression data (15). In this study, we used GSE118307 dataset which includes miR-124 knockout in inducible-neurogenic cell line (iNGN) cells, RNA-seq data (16). It contains a 5-time point for gene expression (69 samples), 0, 1, 2, 3 and 4 days after miR-124 knockout.

### RNA-seq data analysis

We employed Galaxy platform for the analysis of gene expression data. Raw data extracted from GEO into the Galaxy. In the first step, the quality of reads was examined by fastqc. Low-quality read and short read filtered by Cutadapt (minimum length: 30 and law quality: 20), then reads aligned with the reference genome (hg38) using HISAT2. After that, gene count data were generated using FeatureCount. Finally, differentiation expression was performed by DEseq2.

### Transcription Factors finding

Transcription factors (TF) by recognize specific DNA sequences control transcription. Consequently, we investigated which TF is down or upregulated in the miR-124 KO. To reach this goal, we used The Human Transcription Factors database (17).

### Functional Enrichment Analysis

Gene set enrichment analyses (GSEA) are among the most vastly used techniques to evaluate functional gene expression data interpretation. This way, several statistical methods and software were developed (18). GSEA tools summarize genomic data in higher-level biological features. To achieve this aim, GSEA 4.1.0 desktop application was employed for find canonical pathways base on KEGG data (19). FDR < 0.05 was set as the threshold. Gene Ontology (GO) functional enrichment analysis was performed by Enrichr (20).

### Investigation of genes associated with neurodegenerative diseases

The DisGeNET Database includes human gene-disease associations and variant-disease associations information (21). To discover neurodegenerative related genes, we used DisGeNet database. Upregulated and downregulated genes were compared with genes involved in AD, PD, HD and MS, overlap genes were selected. To reach this goal, genes related to AD, PD, HD and MS were downloaded from DisGeNET, then they were compared to upregulated and downregulated genes.

### Protein and genetic interactions

GeneMANIA is a user-friendly database which provides protein and genetic interaction. For this propose, we employed the GeneMANIA to evaluate protein and genetic interaction for upregulated and downregulated genes in all of five-time points (22).

## Result

### Alzheimer’s disease

miR-124 is highly express in the brain; however, it is dysregulated in the hippocampus of AD animals. Downregulation of miR-124 increases Aβ and a variety of cerebromicrovascular impairments and C1q-like protein 3 (C1ql3) alteration. A study on APP/PS1 transgenic mice has shown that miR-124 is involved in angiogenesis and vascular integrity in the hippocampus and cerebral cortex via C1ql3 regulation (23) (Figure 3). Astaxanthin suppresses accumulation of Aβ and inhibit β-site amyloid precursor protein cleaving enzyme 1 (BACE1) as antioxidant, as well as elevating nuclear factor erythroid-2-related factor 2 (Nrf2) and miRNA-124 expression (24). Other non-coding RNAs have a role in AD. lncRNA X-inactive specific transcript (XIST) is upregulated in the in vivo and in vitro AD model. Knockdown of lncRNA XIST elevates miR-124 expression and downregulates BACE1 expression. miR-124s have a direct interaction through lncRNA XIST and BACE1 (mRNA) (25). lncRNA NEAT1 is upregulated in the AD. Knockdown of NEAT1 has protective effects on cellular AD model. Downregulation of NEAT1 upregulates miR-124, so miR-124 can affect Aβ as BACE1 regulator (26). Moreover, it induces miR-124 via adeno-associated virus in APP/PS1-AD mice. It also improves AD-mouse behavior significantly, such as social recognition test and plus-maze discriminative avoidance task (27). Aβ protein (Swedish mutant) affects neuron-microglia cells. When SH-SY5H cells and CHME3 microglia exposure extracellular APP and Aβ accumulation, they express pro-inflammatory and pro-resoling genes. This phenomenon is associated with miR-155, miR-146a and miR-124 secretome from SH-SY5H cells. Microglia internalizes these exosomes and recapitulates the cells of origin. These exosomes are capable of inducing acute and delayed microglial upregulation of TNF-α, HMGB1 and S100B pro-inflammatory markers (28). Thymoquinone is an anti-inflammatory and neuroprotective agent. Studies on Streptozotocin-induced neurodegeneration in the rats have shown that Thymoquinone activates JNK protein, upregulates miR-124 and downregulates ERK1/2 and NOS enzymes (29). miR-124 expression is downregulated via Aβ25-35 in neuronal cells, but upregulates domain death agonist. Domain death agonist is miR-124 target, so miR-124 upregulation reduces Aβ25-35 toxicity (30). Transfect of AD mice (dentate gyrus) via miR-124 lentiviral vector improves biochemical and learning ability. Biochemical alteration includes down-regulation of Bcl-2 to Bax ratio increase in Beclin-1 and decrease in expression LC3II, Atg5 and p62/SQSTMl expression (31). But another study reported that miR-124 overexpression in the hippocampus of P301S transgenic mice downregulate PTPN1 expression and caused hyperphosphorylation of tau. Another study showed that miR-124 increases in temporal cortex and hippocampus of AD patient (32). Another study reported miR-124-3p attenuates cell apoptosis ad Tau hyperphosphorylation and increases expression of Caveolin-1, phosphoinositide 3-kinase (PI3K), phospho-Akt (Akt-Ser473)/Akt, phospho-glycogen synthase kinase-3 beta (GSK-3β-Ser9)/GSK-3β) in N2a/APP695swe cells (33). miR-124 downregulates in aging in BV2 microglia cells. In oxidative stress condition, miR-124 downregulates and RFX1 increases protein level. miR-124 targets RFX1 mRNA and RFX1 bind in the first intron of ApoE gene (34). miR-124 targets 3’UTR Notch ligand Delta. Reduction of Delta expression extended lifespan and meliorated learning defects of AD fly fruit (35). Resveratrol is a natural compound which have neuroprotective role in AD. However, Resveratrol downregulates miR-124 and miR-134 (36). Melatonin is the crucial hormone for the regulation of the circadian rhythm and it decreases in the AD patients. Inhibition of melatonin induces AD-like memory deficits. Melatonin could downregulate miR-124 by promoting EPAC expression (37). Moreover, miR-124 is involved in APP splicing (38).

**Figure 3.**
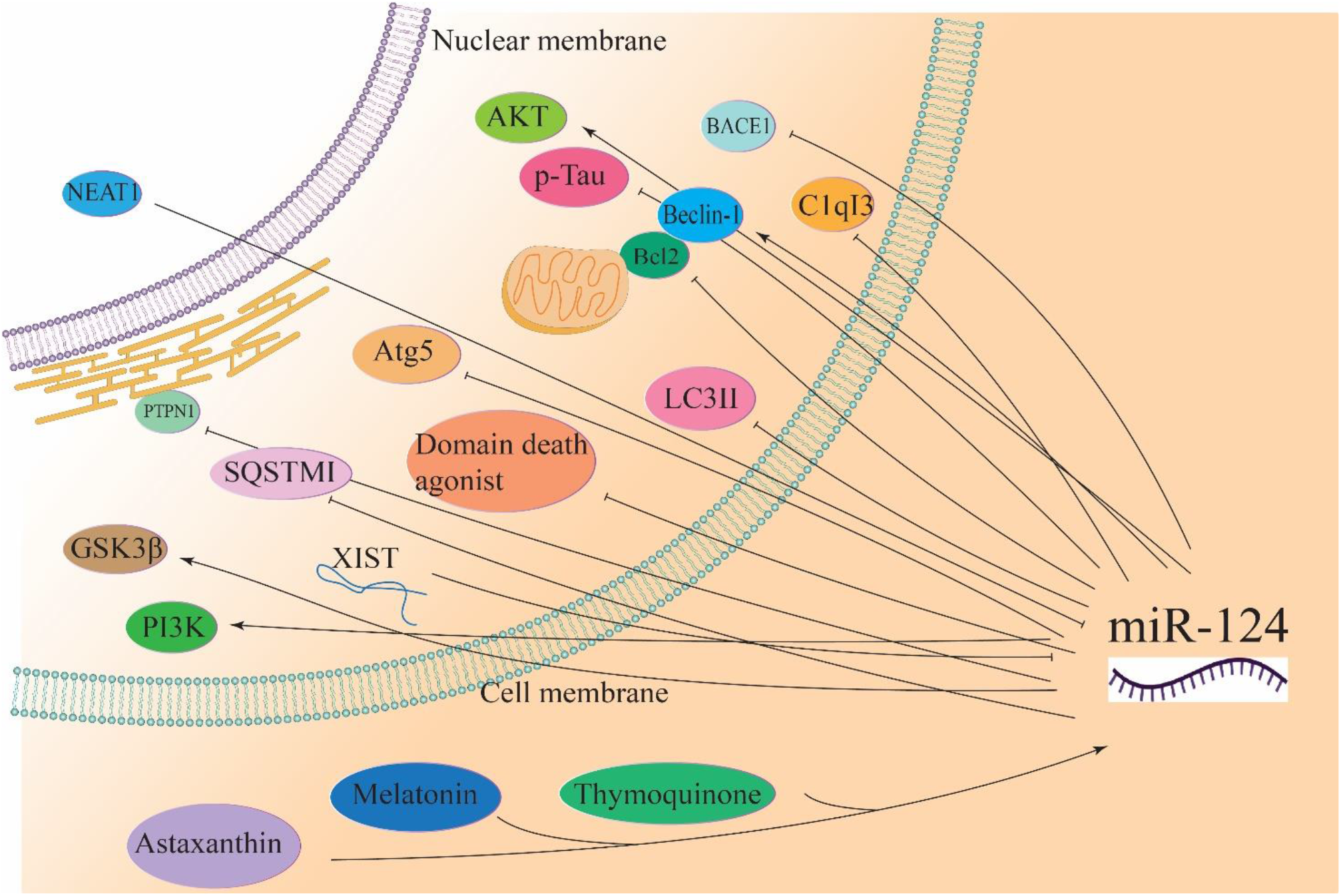
miR-124 function on AD.

### Parkinson’s disease

HOXA11-AS long non-coding RNA mediated neuronal injury in MPTP cellular model of PD and LPS-induce microglia activation, but miR-124 has opposite effect. HOXA11-AS over-expression increases expression of inflammatory factor and FSTL1, NF-kB and NLRP3 inflammasome while miR-124 is the target of HOXA11-AS and FSTL1 (39). KLF4 and NEAT1 are upregulate in the midbrain of MTPT-HCl mice. Knockdown of NEAT1 decreases apoptosis and increases cell viability in MPP+-treated SH-SY5Y cells while upregulation of KLF4 could reverse this process. miR-124 targets both of NEAT1 and KLF4. Downregulation of NEAT1 promotes cell viability and decreases apoptosis via miR-124 in MPP+-treated SH-SY5Y cells (40). miR-124 is upregulated in PD blood plasma (41). KPNB1, KPNA3, and KPNA4 upregulate in the PD brain. KPNB1, KPNA3, and KPNA4 regulate NF-κB/p65 nucleo-cytoplasmic transport. KPNB1, KPNA3, and KPNA4 are miR-124 target’s genes (42). DAPK1 is involved in gamma-interferon induced programmed cell death. Overexpression of DAPK1 in the dopaminergic neurons, injury’s this neurons and locomotor disabilities in the PD mice. DAPK1 negative correlates with miR-124 in SH-SY5Y cells treated by MPP+. miR-124-3p mimic significantly inhibits DAPK1 expression and attenuates MPP+-induced cell. MALAT1 knockdown improves behavior and decreases apoptosis by DAPK1 downregulating and miR-124 upregulating (43). MALAT1 increases apoptosis by sponging the miR-124 (44). miR-124 promotes dopamine receptor expression and neuronal proliferation. Moreover, miR-124 suppresses neuronal apoptosis via downregulating EDN2 and Hedgehog signaling activation (45). miR-124 transfection has shown that miR-124 targets MEKK3 and NF-kB (46). miR-124 also targets STAT3 as inflammation gene (47). miR-124 targets ANXA5 which is associated with the simulation of the ERK pathway (48). miR-124 increases the migration of the new neurons into the 6-OHDA PD model striatum (49). miR-124 is downregulated in PD plasma but miR-124 increases in PD with depression to PD without depression (50). miR-124 downregulates in cellular MPTP model and this downregulation increases apoptosis and increases autophagy through Beclin1 an increase in LC3 II/LC3 I. miR-124 inhibition decreases p-mTOR and increases p-AMPK. miR-124 protects dopamine neuron in PD through apoptosis and AMPK/mTOR regulation (51). It has been demonstrated that miR-124 elevates PD brain repair in subventricular zone. miR-124 induces migration of neurons into the lesion stratum in the PD mice model (52). Bim, which is another target gene for miR-124, inhibits Bax translocation to mitochondria (53).

**Figure 4.**
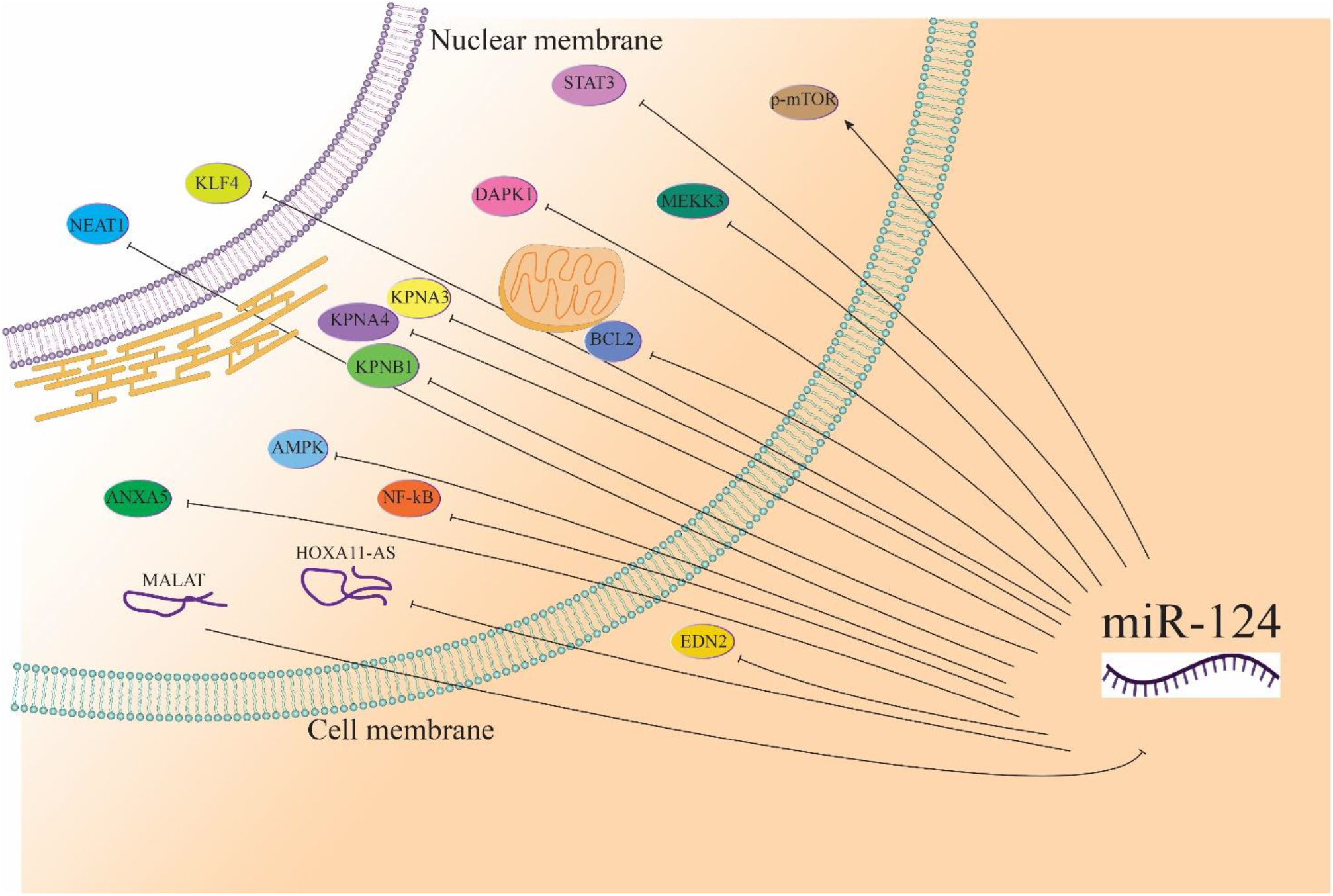
miR-124 function on PD.

### Multiple sclerosis

miR-124 is decreased in the MS mice model. However, anti-inflammatory drugs, such as Bu Shen Yi Sui capsule (is a traditional Chinese medicine) decreases inflammation and increases miR-124 level (54). miR-124 is an anti-inflammatory marker which strongly decreases in MS patient, especially in progressive MS (55). Resveratrol is another natural compound which decreased severity of MS, including inflammation and central nervous system immune cell infiltration. Resveratrol significantly upregulates miR-124 and downregulates SK1 miR-124 target gene. Resveratrol improves amelioration of MS model and suppresses inflammation via alteration of miR-124/SK1, thereby it inhibits promotion of apoptosis (56). miR-124 increases in MS patient blood compare to control (57). However, miR-124 increases in the demyelinated postmortem MS hippocampi. AMPA glutamate receptors, namely GRIA1, GRIA2 and GRIA3, are miR-124 target genes, and these genes downregulate in MS (58).

### Amyotrophic lateral sclerosis

miR-124-3p level is decreased in the spinal neuron of ALS mouse models. Increase of miR-124-3p is associated with spinal motor neuron-derived exosomes in ALS mice (59). miR-124 influences on astrocytic differentiation. It has been demonstrated that miR-124 targets Sox2 and Sox9 in the ALS transgenic mice (59).

### Huntington’s disease

miR-124 is repressed in HD. Injection of miR-124-exosome into R6/2 transgenic HD mice striatum decreases REST expression (60). Studies have shown that miR-124 is downregulated in HD transgenic cells and animal model. miR-124 targets CCNA2. CCNA2 is involved in cell cycle process and knockdown of CCNA2 alters proportion of cells in S phase of HD ell model (61).

**Figure 5.**
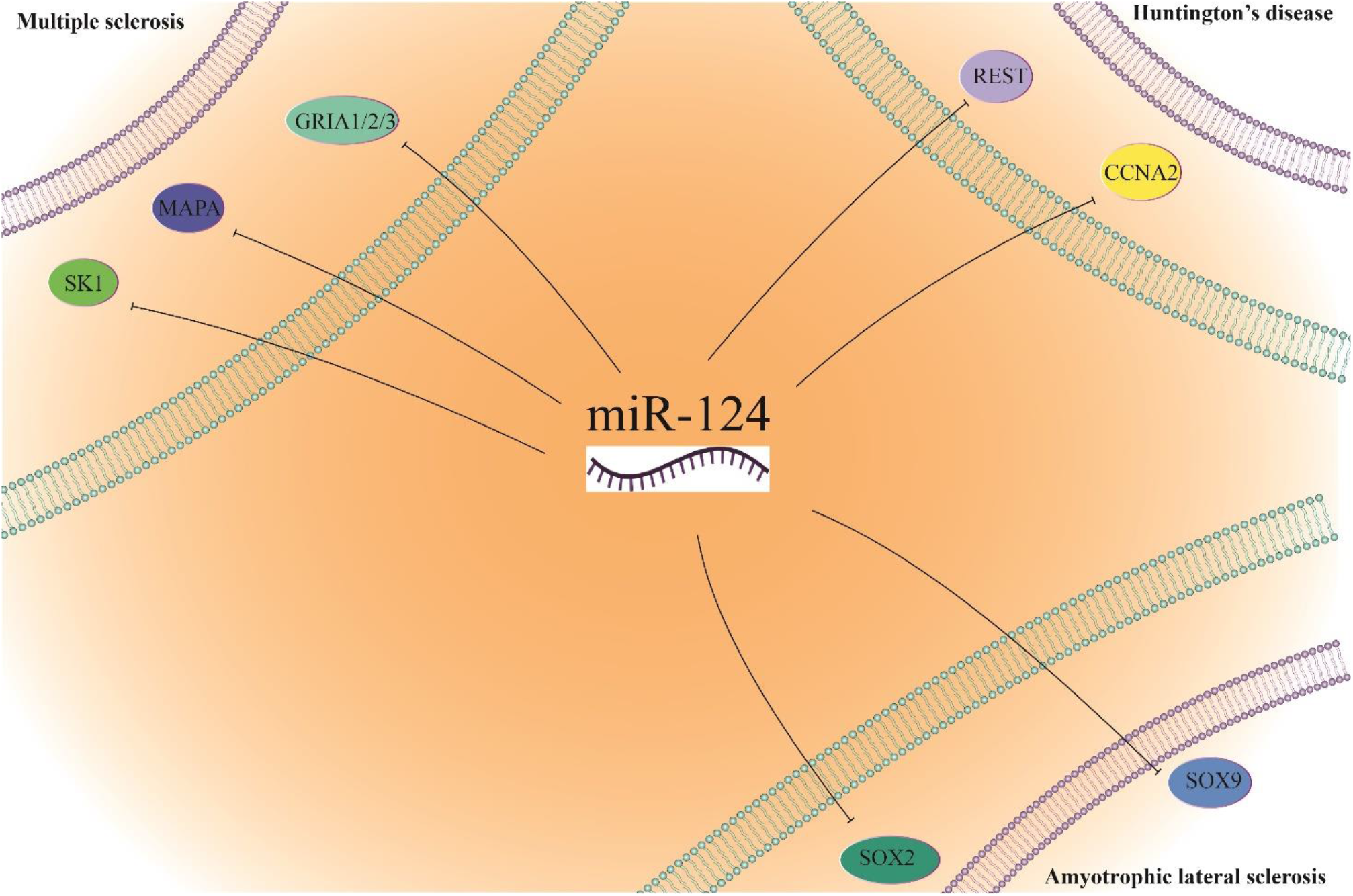
miR-124 function on MS, HD and ALS.

Both mature type of miR-124 (3p and 5p) is highly express in the neuron system (12).

### Differentially expressed genes identification

We investigated the effect of knockdown miR-124 on the neuronal cell in 5 time point (Figure 6). To achieve this goal, RNA-seq data was extracted from GEO and evaluated gene expression differentiation, which included up and downregulated gene in this five-time points. miRNA’s role is post-translational regulation of genes and it inhibits translation their targets, which is why in this study, we evaluated downregulation of genes too.

**Figure 6.**
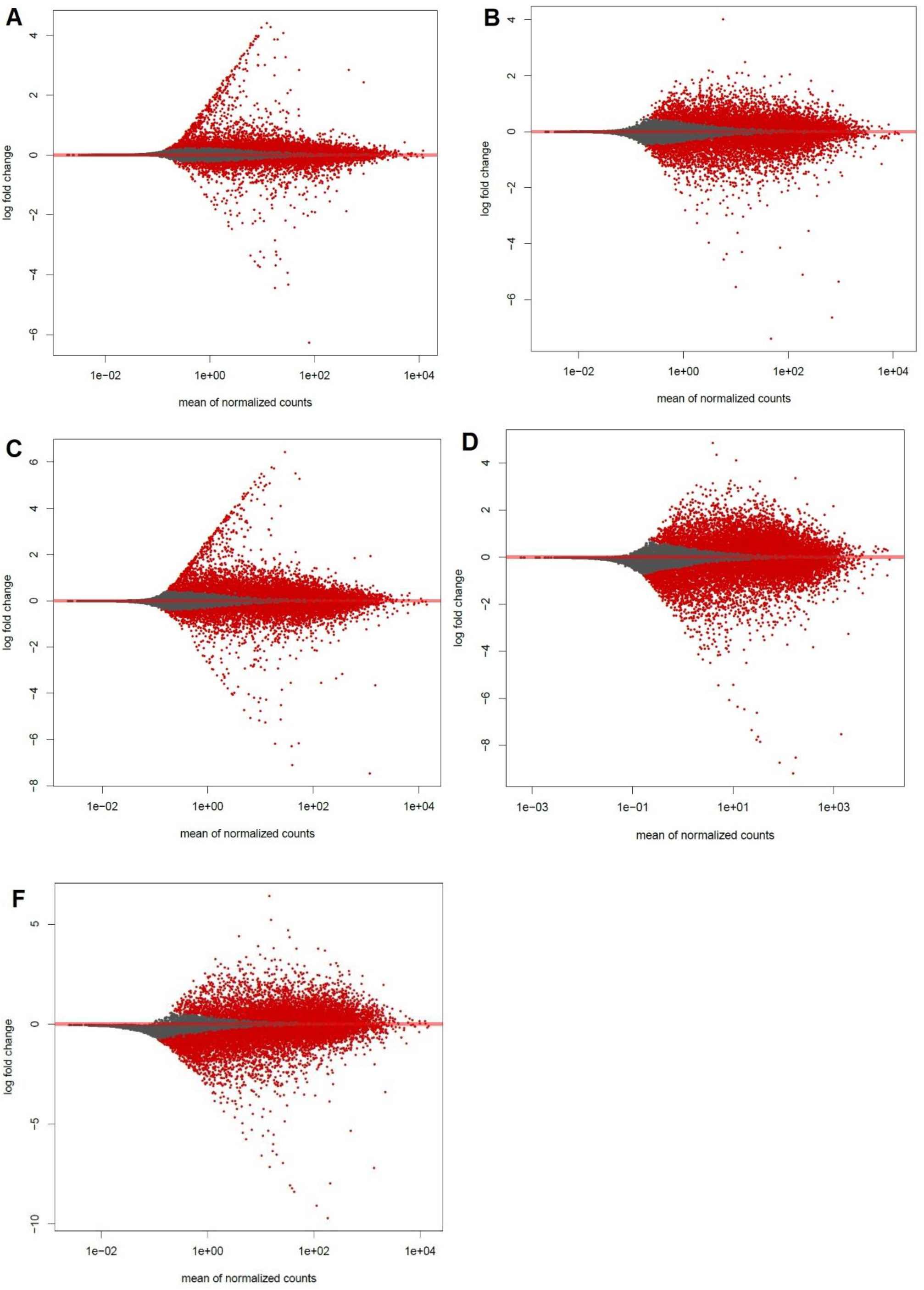
The pattern of differential expression genes after Knockdown miR-124. In order **A)** day 0, **B)** day 1, **C)** day 2, **D)** day 3, **F)** day 4.

On the day 0 and day 2, FLG is the highest expression (Table 1). On the days 1,3 and 4 MGAT4C, CCL1 and PADI2 were the highest expression respectively. Chemokine CCL1 is express in APP/PS1 mice but that is higher in wild types. CCL1 is involved in immune cells by CCR8 receptor interaction (62). In the day 0, 1 and 2, PEG3 was the lowest expression. ANKRD45 is the lowest expression in the day 3 and 4. ANKRD45 have a dynamic role in mitosis and knockout of this gene induce apoptosis in cell culture (63).

**Table 1.** Top five up and downregulated genes (sorted by foldchange).

| Time point | Upregulated genes | Downregulated genes |
| --- | --- | --- |
| **Day 0** | FLG<br>TLDC2<br>TMEM140<br>SLC4A5<br>BEND3P3 | PEG3<br>PRSS57<br>EMC6<br>COX20<br>HYI |
| **Day 1** | MGAT4C<br>C1orf94<br>GDPD2<br>MYH2<br>EGR1 | PEG3<br>AJAP1<br>ABCA5<br>MAFA<br>PRSS57 |
| **Day 2** | FLG<br>RGL4<br>C22orf23<br>BEND3P3<br>TMEM140 | PEG3<br>ANKRD45<br>AJAP1<br>TMEM200B<br>HYI |
| **Day 3** | CCL1<br>CD248<br>WNT11<br>IGFBP4<br>C10orf54 | ANKRD45<br>TTLL10<br>PEG3<br>ZNF662<br>CAV1 |
| **Day 4** | PADI2<br>CCL1<br>SRL<br>SSTR5<br>POU3F4 | ANKRD45<br>PEG3<br>TTLL10<br>AJAP1<br>PRO1804 |

Upregulation genes varied on different days, but the decrease in gene expression has increased over time (Figure 7). It seems when miR-124 expression occurs, it activates many genes and it has an activation role instead of repression one. It seems that miR-124 has an active role in neurons and causes upregulation many genes.

**Figure 7.**
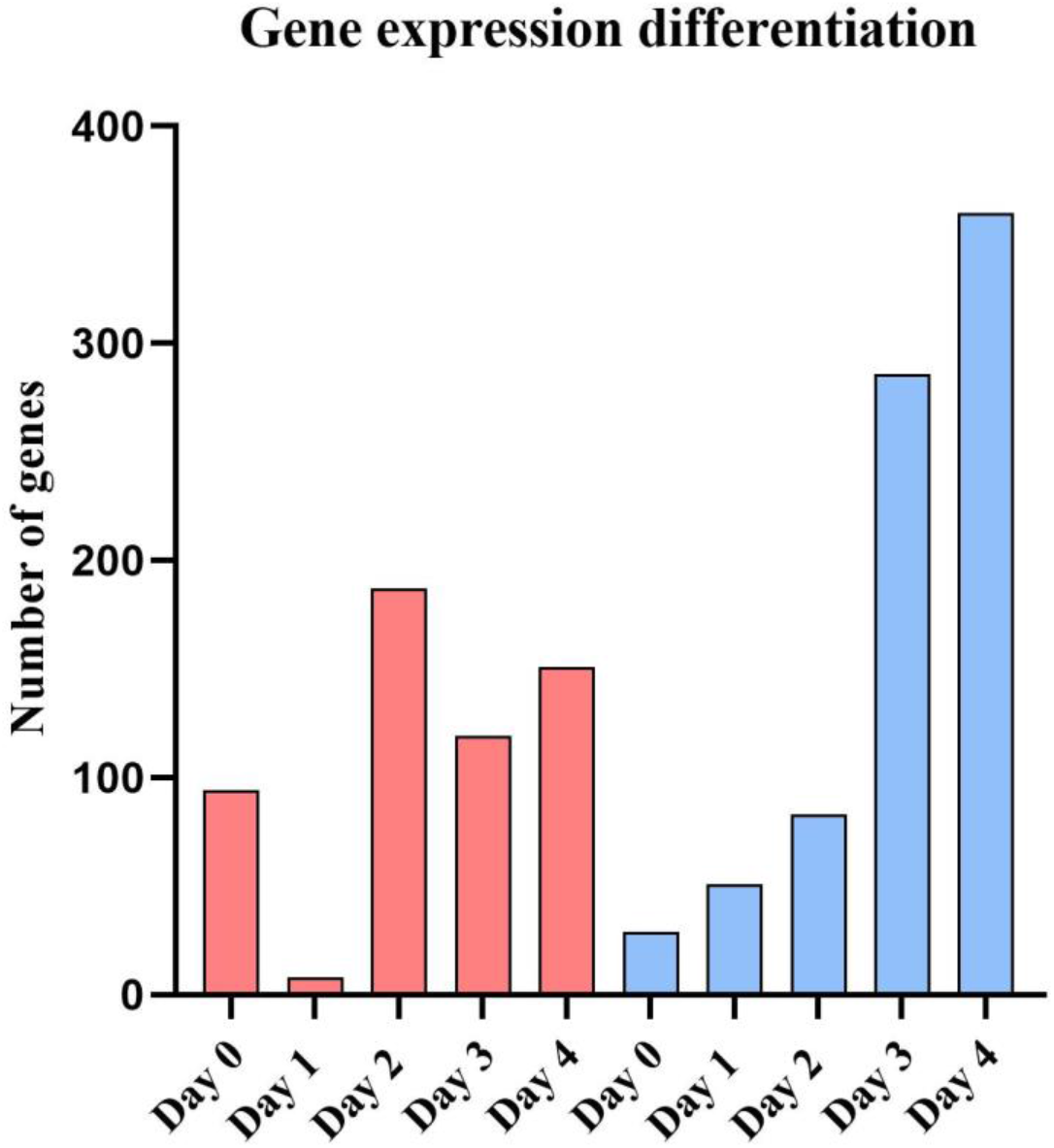
Figures related to up (red bars) and downregulated (blue bars) gene by miR-124 knockout.

In the upregulated gene was not any common genes, but 7 genes were common in the downregulated genes (Figure 8). These genes include: paternally expressed 3 (PEG3), adherens junctions associated protein 1 (AJAP1), cathepsin F (CTSF), serine protease 57 (PRSS57), zinc finger protein 662 (ZNF662), pyridoxamine 5’-phosphate oxidase (PNPO) and RNA binding motif protein 46 (RBM46). Inactivation expression of PEG3 induces deficits in olfactory function, sexual and maternal behaviors, oxytocin neuron number, metabolic homeostasis and growth. Female Peg3^+/-^ mice have reduced the number of oxytocin neurons and males Peg3^+/-^ have reduced neuronal activation (64). AJAP1 is expressed mainly in neuronal cells and it is associated with γ-Aminobutyric acid type B receptor subunit 1. AJAP1 is involved in epilepsy. Overexpression of AJAP1 can reduce the frequency of spontaneous seizer (65). CTSF is a member of lysosomal cysteine protease of the papain family (66). Mutation in CTSF causes type B Kufs disease, which is a type of ND. Mutation of CTCF is also identified in AD patients (67). PRSS57 is involved in innate immune reactions as regulatory properties towards neutrophil responses. PRSS57 silencing increases mitochondria activity in the neurons and it seems that inhibition of PRSS57 promote CD34^+^ cell proliferation in the MS (68). RBM46 plays a role in stem cell differentiation. It regulates the degradation of β-Catenin mRNA. RBM46 (69).

**Figure 8.**
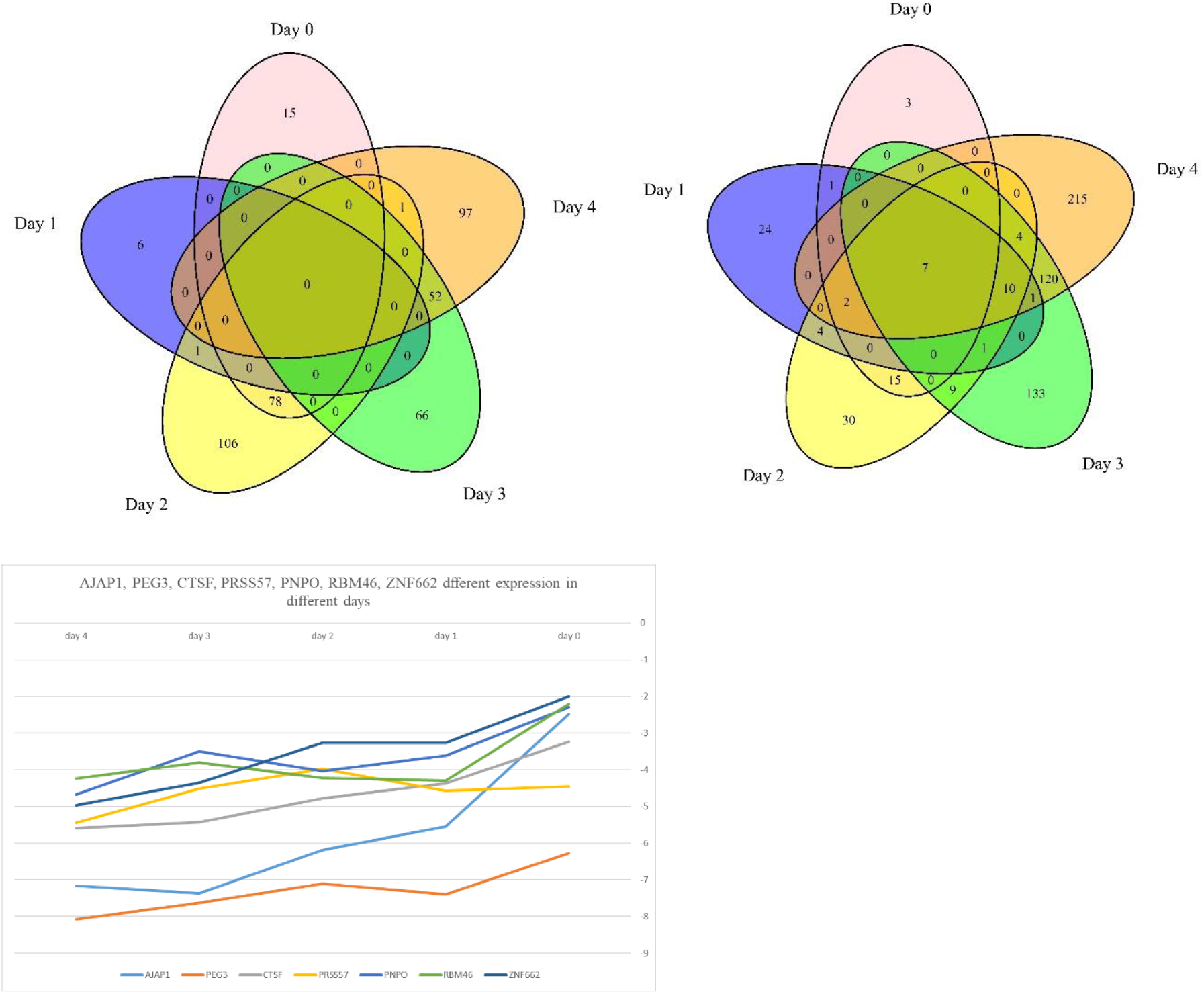
Common genes in up and downregulated gene via miR-124 knockout.

### Transcription factors expression

We evaluated which transcription factor is altered during downregulating miR-124. 72 TFs were altered during 5 days. The most TF differential expression was in the day 4. In the first day’s TF alteration expression, there was not much change. However, over the time, TF alteration expression is increased. Zinc finger protein 367 (ZNF367) and paternally expressed gene 3 (Peg3) were downregulated from first day to fifth day. ZNF367 was a TF involved in kinesin family member 15 (KIF15). It has been demonstrated that ZNF367 silencing represses cell growth of breast cancer (69).

We evaluated which upregulated TF could directly be targeted by miR-124. In this context, miR-124 and upregulated TF interaction were verified by TargetScan. We found seven TF, including NFIB, CDX2, DLX2, ELK3, NFIX, NR4A1, TAL1 could direct target to miR-124.

### Evaluation relationship differential expression gene form miR-124 KO to AD, PD, MS, HD

Database DisGeNET was used to investigate the association between changes in gene expression caused by miR-124 KO and neurodegenerative diseases. During these 4 days and an increase in differential expression genes, neurodegenerative related genes were increased.

On day 0 PINK1 and day 1 EGR1 are multi-disease genes for upregulated genes. In downregulated genes, SLC6A4 is a multi-disease gene at day 1. In addition of EGR1, IL7 is multi-disease on day 2 for upregulation and SLC6A4 and CLDN5 are for downregulation genes. At day 3, the number of multidisease genes increased. CALB2, SLC2A4, TNFRSF1B and NOS1 are multi-disease for upregulation and MIR200A, CAV1, SLC6A4 and ADCYAP1 for downregulated genes. On day 4, have most multi-disease genes. On this day, NOS1, BCHE, TNFRSF1B were for upregulated genes and GJA1, TAC1, MIR200A, SLC40A1, TAC1, SPP1, CAV1, LPAR3 and ADCYAP1 were for downregulated genes multi-disease genes.

**Figure 9.**
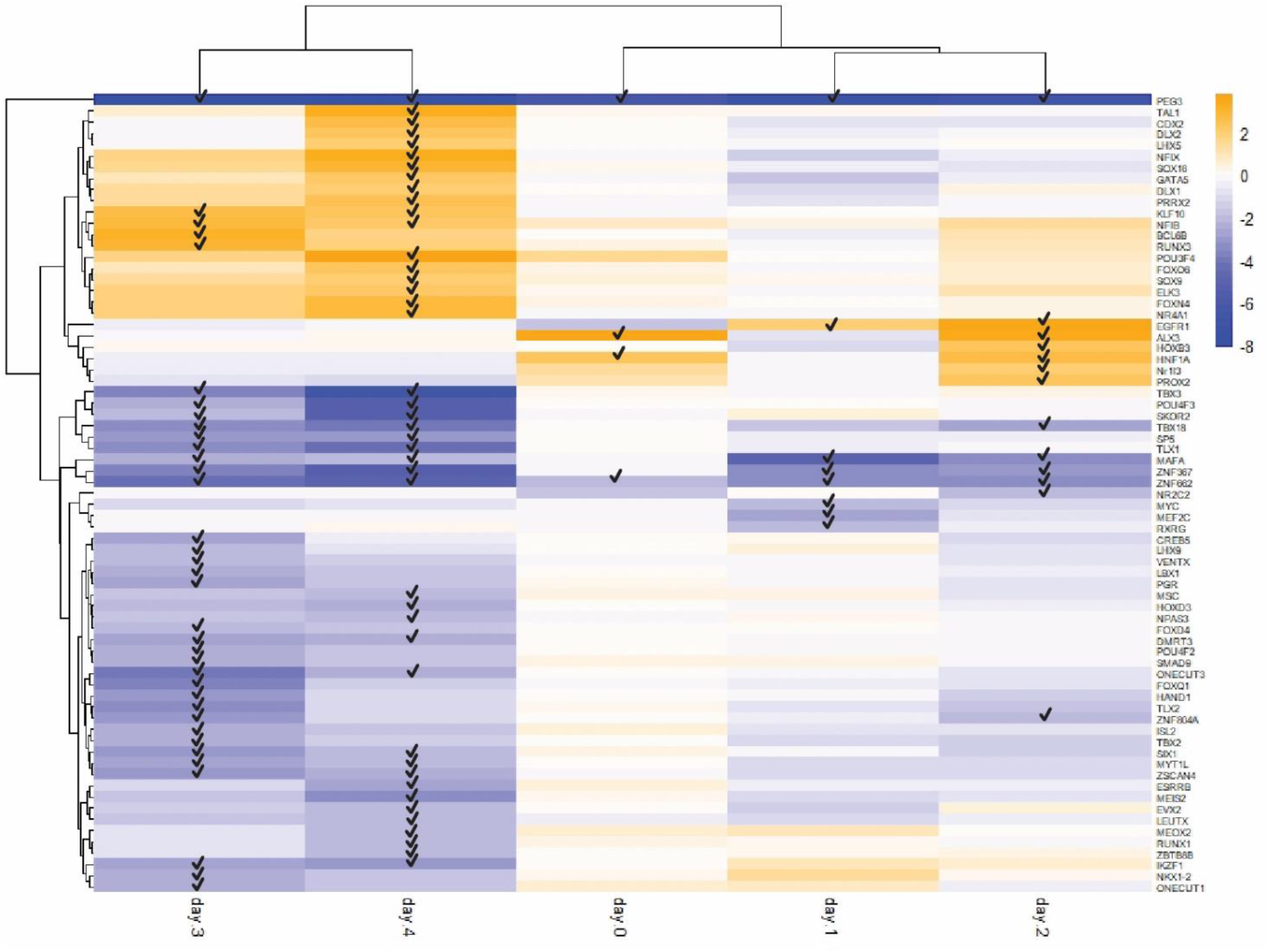
Heatmap form TFs expression. Each tick means, FDR < 0.05.

**Figure 10.**
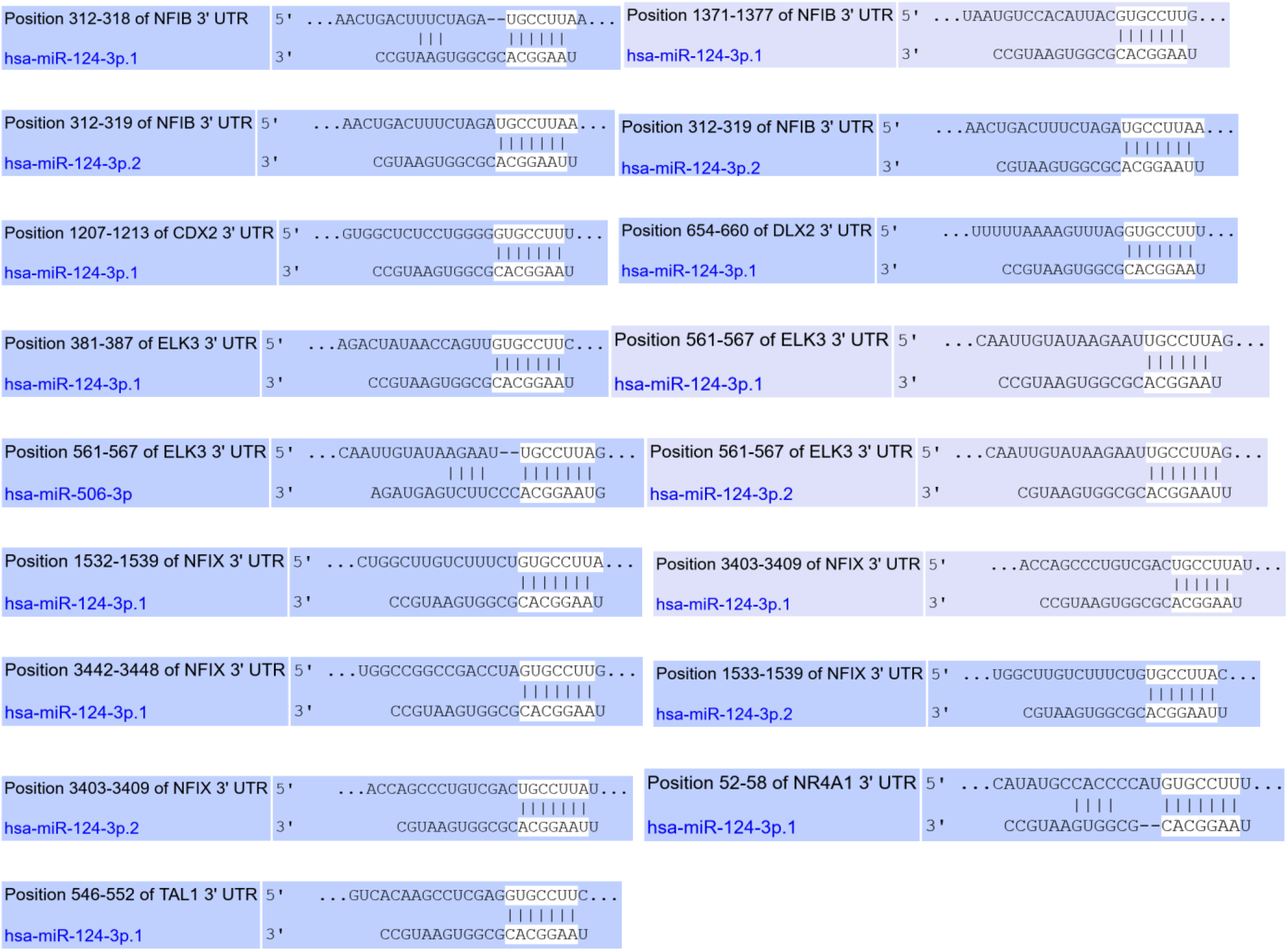
Interaction of miR-124 was done by upregulated TF. Several TF upregulated after miR-124 inhibition, which is why the interaction of miR-124 with upregulated TFs were investigated. 7 TFs were identified that could be targets of miR-124.

**Figure 11.**
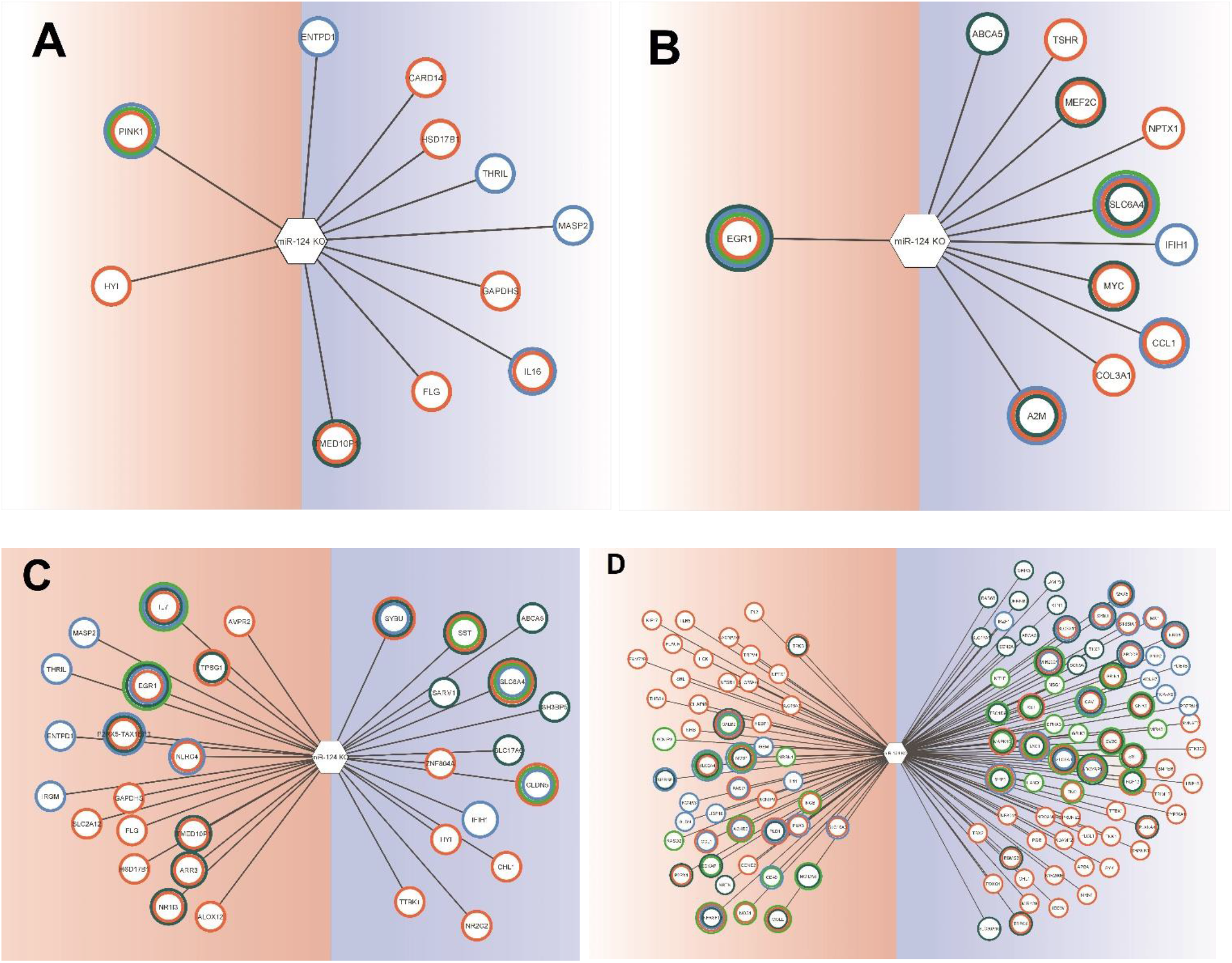
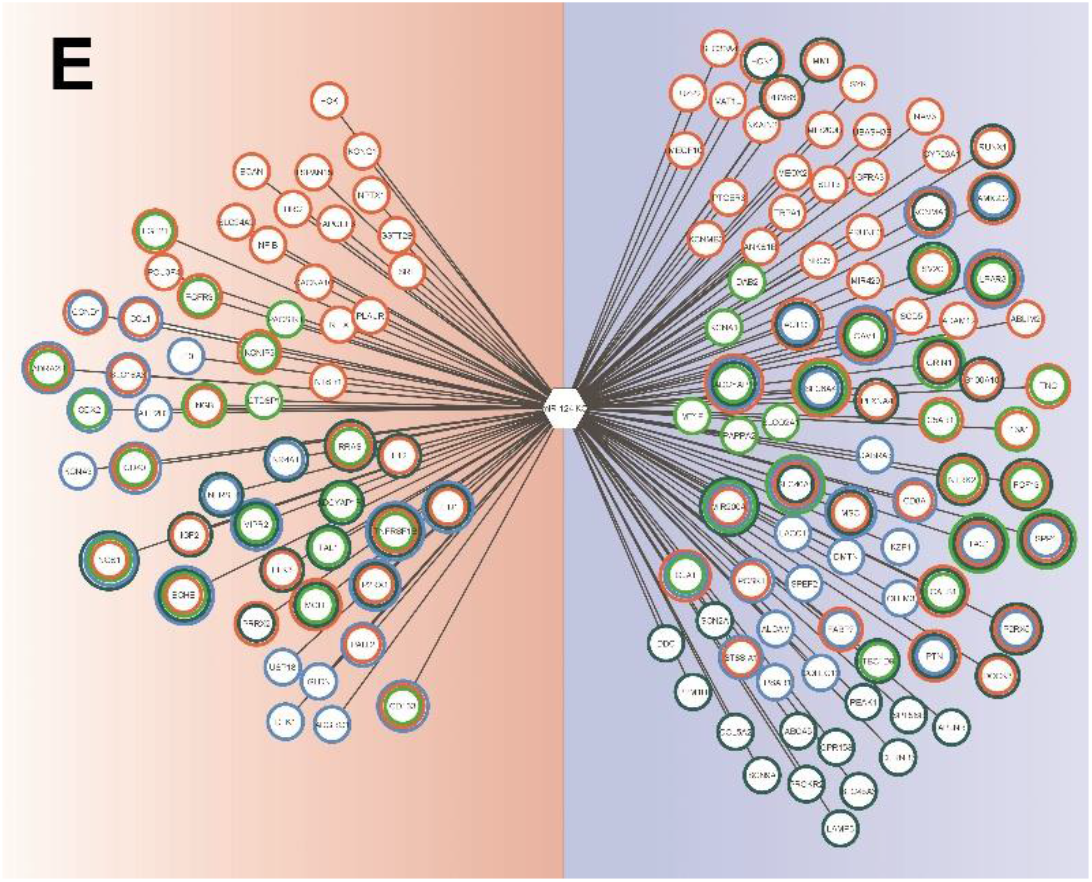
Up and downregulated gene were contributed by ND. **A)** day 0 **B)** day 1 **C)** day 2 **D)** day 3 and **E)** day 4. Red area contributes to upregulated genes and blue area contributes to downregulated genes. Orange border indicates AD, light green shows PD, blue displays MS and dark green illustrates HD.

### Gene ontology

The enriched categories plotted in chord diagram. Each diagram contains 2 part GO terms and related genes. GO terms divided into three areas, which include biological process, cellular component and molecular function. These three areas were highlighted by purple, orange and green colors. In each plot, we used top seven term (base on p-value).

**Figure 12.**
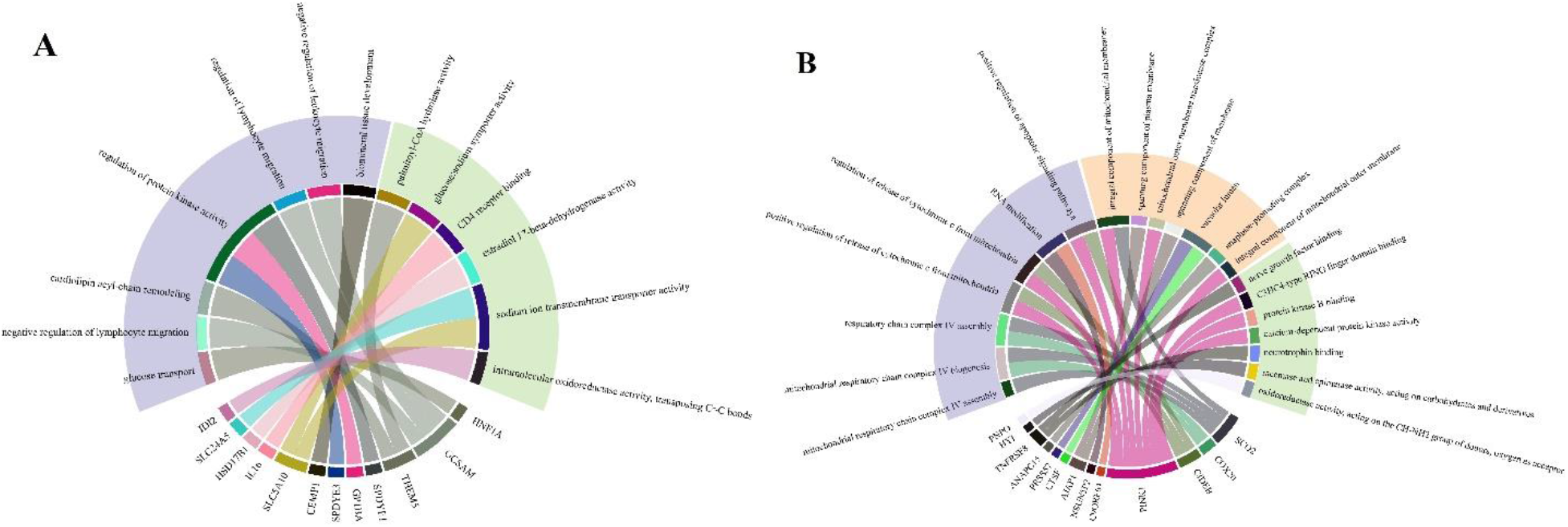
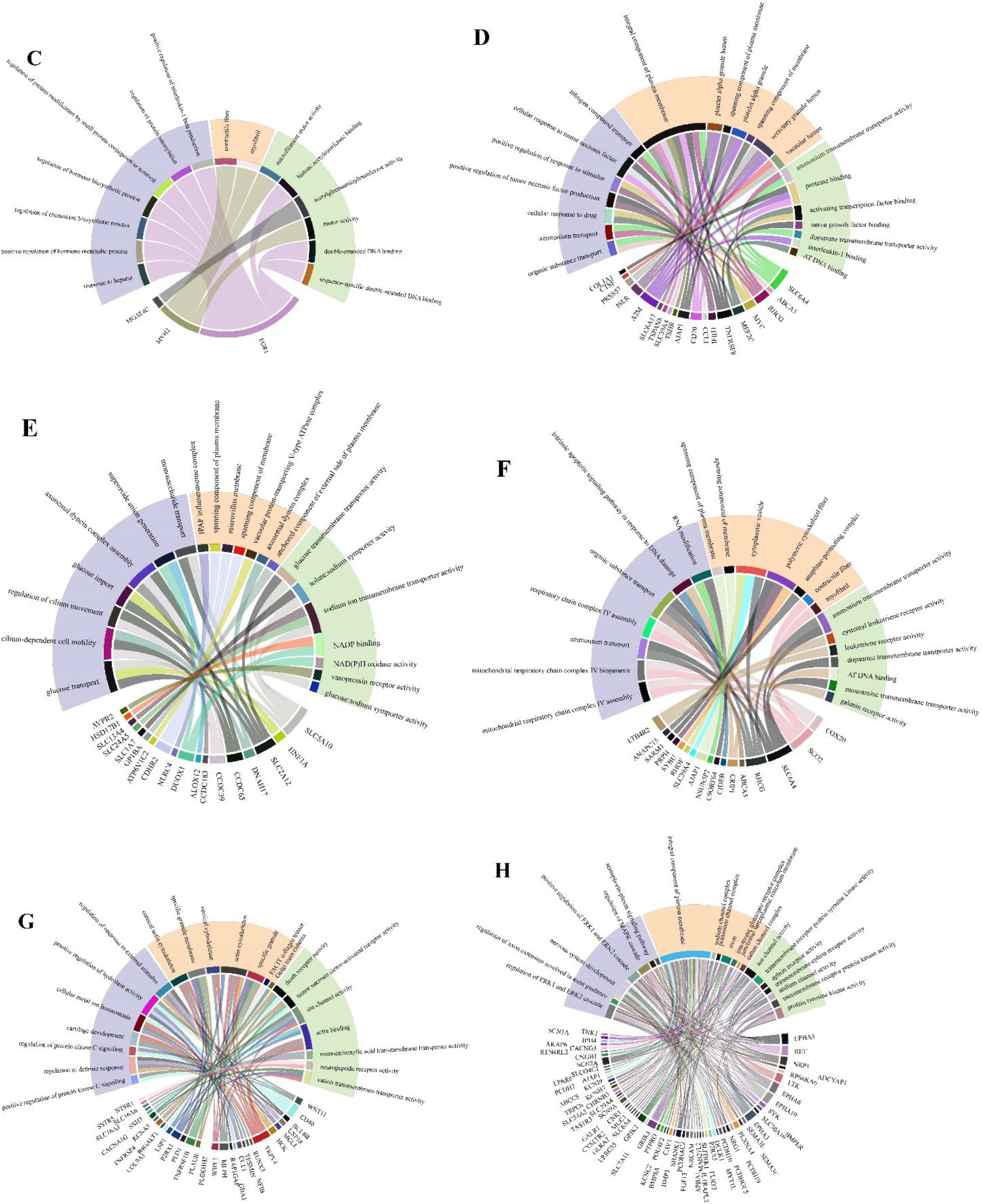
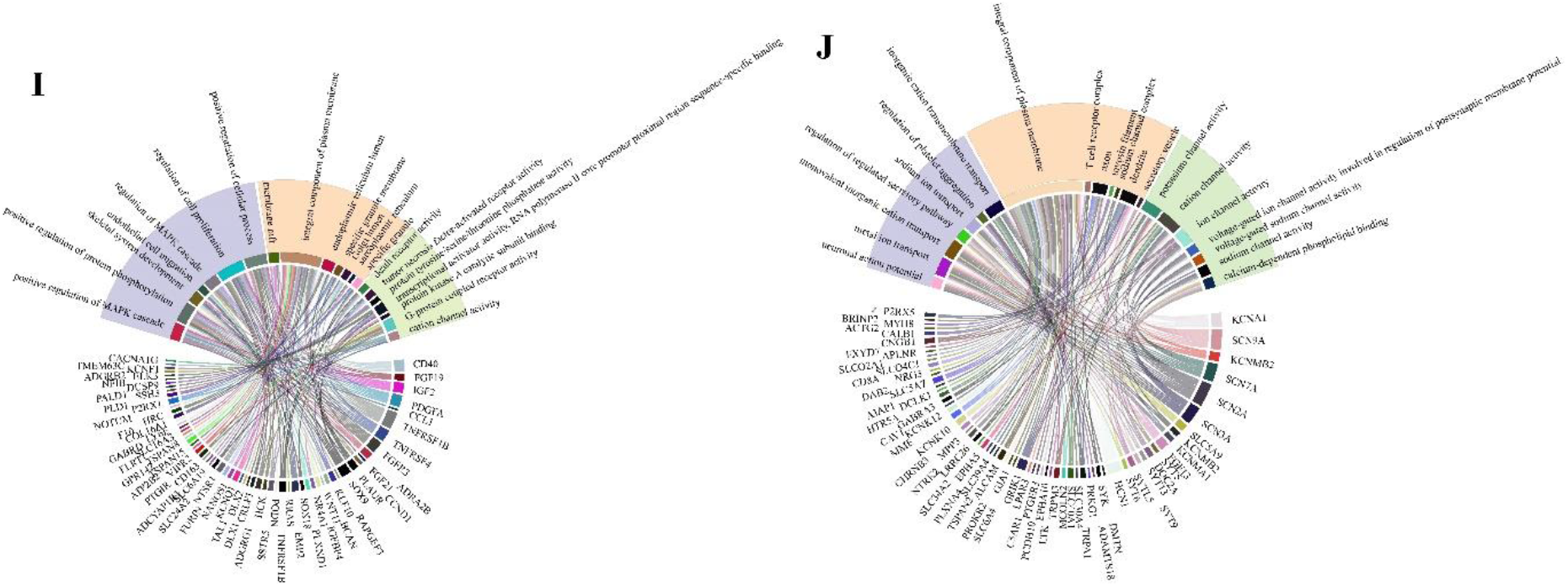
Result of GO analysis. A, C, E, G and are related to GO analysis of upregulated genes on day 0, 1, 2, 3 and 4. B, D, F, G and h are related to GO analysis of downregulation genes on day 0, 1, 2, 3 and 4.

### Gene network

Gene-gene interaction was performed by GenMANIA Cytoscape plugin. Gene network was plotted, then it was analyzed by cytoscape. TM6SF1, EGR1, FND38, FGR, ITPKB, NFIB (three genes have the same degree for fourth day) and CSF3R have the most interaction on days 0, 1, 2, 3 and 4 of the upregulation of genes respectively. ANAPC15, MRLP34, EPHX2, CIDEB (four genes have the same degree for 0 day), COL3A1, VIT, CTNNA2 and NTRK2 have the most interaction on days 0, 1, 2, 3 and 4 of the downregulation of genes respectively.

**Figure 13.**
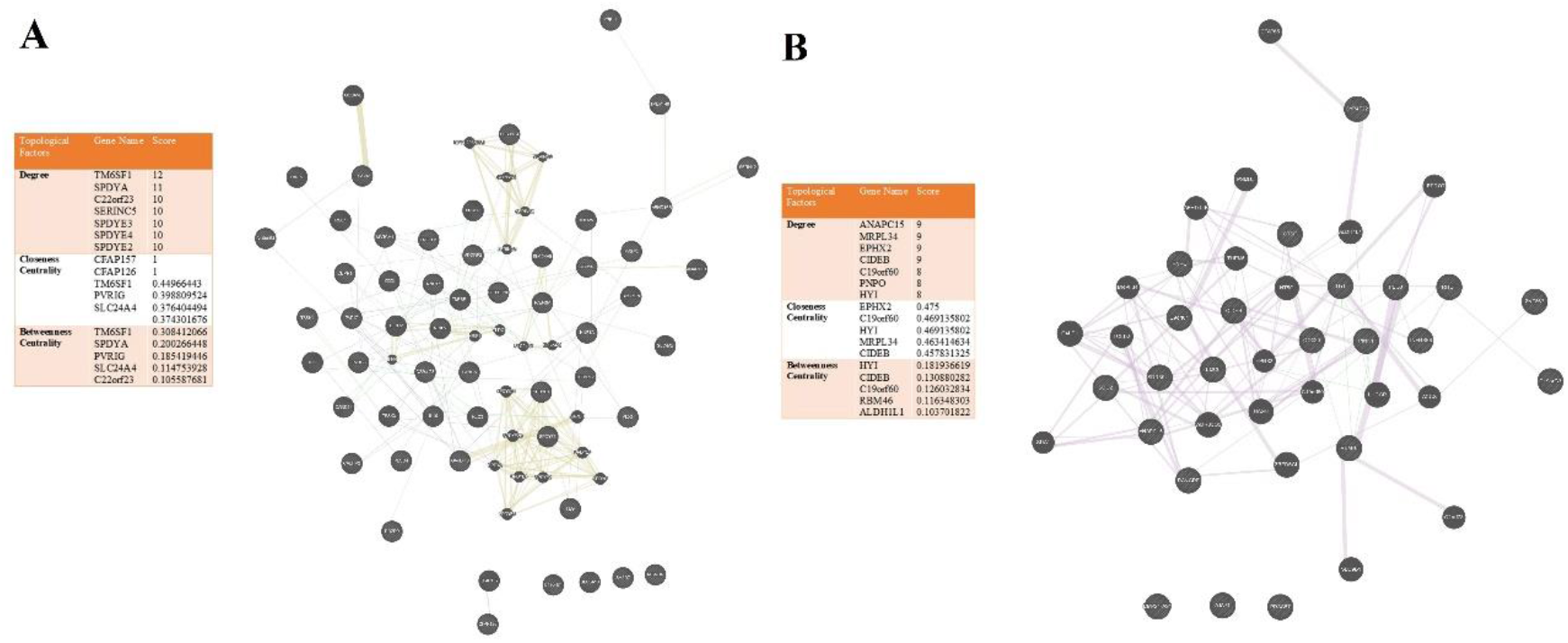
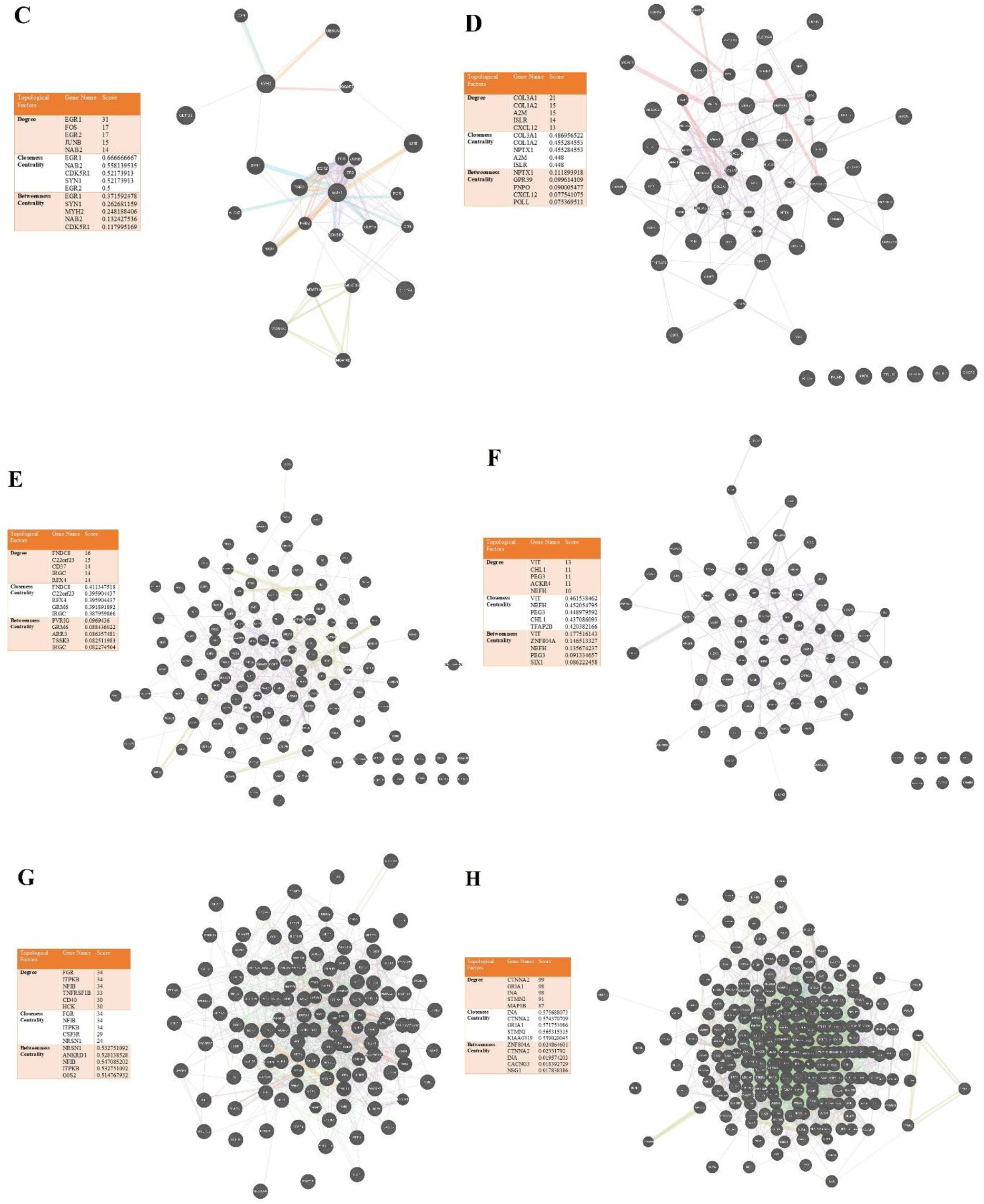
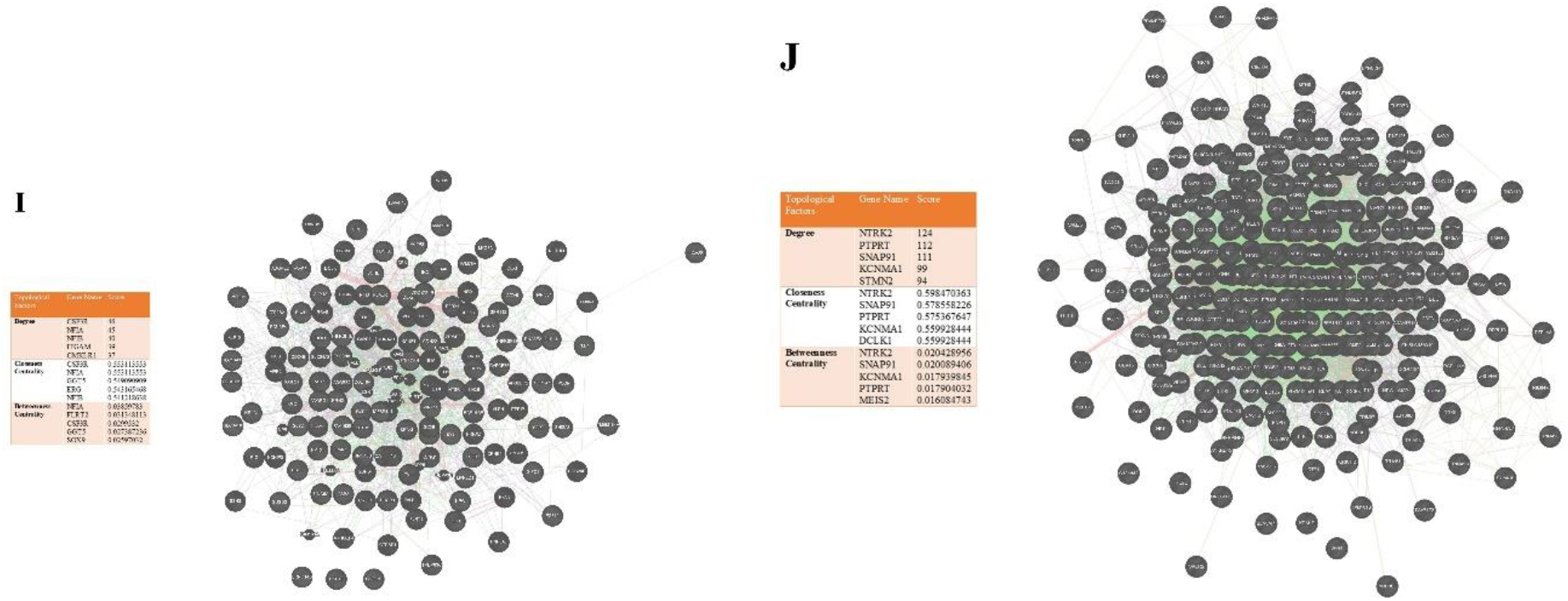
Gene network.

### Pathway analysis

Pathway analysis was performed by GSEA. FDR < 0.05 was considered as statistical threshold according to that, no pathway was obtained for downregulation of genes on the third, fourth and fifth days, as well as upregulation on third day. Result of GSEA was plotted in 3Dplot. X axis illustrates the number of genes (size) for each pathway, Y axis shows the value of FDR, Z axis displays the score of enrichment analysis. In the downregulated gene on day 0, UBIQUITIN_MEDIATED_PROTEOLYSIS and SPLICEOSOME have largest size and, in the upregulation, genes are NEUROACTIVE_LIGAND_RECEPTOR_INTERACTION and MAPK_SIGNALING_PATHWAY. On day 1 ENDOCYTOSIS and JAK_STAT_SIGNALING_PATHWAY have the largest size in downregulation of genes and TIGHT_JUNCTION and SPLICEOSOME for the upregulation of genes. Also, CYTOKINE_CYTOKINE_RECEPTOR_INTERACTION and PURINE_METABOLISM have the largest size for the upregulation of genes on day 3 and CYTOKINE_CYTOKINE_RECEPTOR_INTERACTION and REGULATION_OF_ACTIN_CYTOSKELETON for day 4.

## Discussion

Recently, there are no significant neuroprotective treatment that can effectively improve the diseases, namely AD or PD (70). Genetic factors are major risk for developing ND, but environmental factors also impact the onset progression of ND (71). In the year 2019, nearly 50 million individuals worldwide had neurodegenerative diseases and it is estimated that this figure will rise to 150 million by 2060 (72).

Dysregulation of miRNAs are commonly occurred during ND and that is established by animal model and ND patients. One of the most dysregulations occur on miR-124-3p (73).

In this study, we evaluated the effect of miR-124 on ND by literature review and in silico analysis. According to literature review, it has a functional role in AD. miR-124 inhibits BACE1 (25) which is required for Aβ generation. BACE1 is an important drug target to attenuate Aβ production in early AD (74). Moreover, miR-124 increases Nrf2 expression (24). Nrf2 is a cellular oxidative response. Nrf2 induction AD mice model not only reduces oxidative stress and inflammation but it also improves cognitive impairment and improves learning (31, 75). Hyperphosphorylation of Tau is another important pathogenesis of AD. Hyperphosphorylation of Tau increases accumulation of Tau in neurons and lead to neurofibrillary tangles (76). miR-124 attenuates the cell apoptosis via reduction of Tau hyperphosphorylation (33). There are several conflicting reports on the performance of miR-124. For example, it has been reported that miR-124 increases the cortex of AD and hyperphosphorylation of Tau (32). However, almost all reports indicate that reduced miR-124 expression has beneficial effects on AD.

KLF4 is crucial for oxidative stress and cell death via MPP+. It is increased by MPP+. KLF4 overexpression increases MPP+ neurotoxicity and cell death (77). Through targeting KLF4 directly, miR-124 increases viability and decreases apoptosis MPP+-treated SH-SY5Y cells (40). Inflammation is a common pathogenesis in the PD and is involved in onset or progression of PD (78). miR-124 regulates NF-kB pathway by targeting KPNB1, KPNA3, and KPNA4 as NF-kB/p65 regulator (42). Moreover, miR-124 targets NF-kB and STAT3 (43, 47). DAPK1 is involved in gamma-interferon induced programmed cell death and overexpression of DAPK1 injuries to dopaminergic neurons in PD. However, miR-124-3p mimic inhibits DAPK1 (43).

RNA-seq data showed that gene expression alteration is increased over time by knockouting the miR-124 in the neurons (Figure 7). 7 genes were included, namely PEG3, AJAP1, CTSF, PRSS57, ZNF662, PNPO and RBM46, all of which were common in the downregulated genes (Figure 8). All of these genes are involved in CNS function such as GABA regulation. PEG3 regulates oxytocin receptor expression by directly binding to a genomic area inside the third exon of oxytocin (Figure 15) (79). The oxytocin concentration in the hippocampus and temporal cortex of the AD brain increases while elevated levels of oxytocin in the hippocampus can cause memory impairment associated with AD (80). AJAP1 is associated with β-amyloid precursor protein (APP) complex formation. AJAP1, APP and PIANP bind to the GB1a subunit of GABAB receptors. By stabilizing APP at the cell surface, GB1a protects APP from BACE1-dependent endosomal processing. Cultured hippocampal neurons of GB1a/ mice showed that a 40% increase in secreted A levels when compared to WT littermate mice. (81). CTSF mutation is involved in neurological diseases like frontotemporal dementia and AD. Exome sequencing in a consanguineous family with early-onset AD and frontotemporal dementia reveals a homozygous CTSF mutation (67, 82). PRSS57 expression is increased in the brain of MS rats. Silencing of PRSS57 attenuates inflammation and spinal cord injury from MS rats. Moreover, PRSS57 promotes CD34^+^ cell proliferation in these rats (68). pnpo-/- zebra fish induces seizure and could kill them after 20 days. pnpo-/- decreases neurotransmission, including glutamate, γ-aminobutyric acid (GABA) and glycine (83). RBM46 plays a role in stem cell differentiation. It regulates the degradation of β-Catenin mRNA. RBM46 (69).

**Figure 14.**
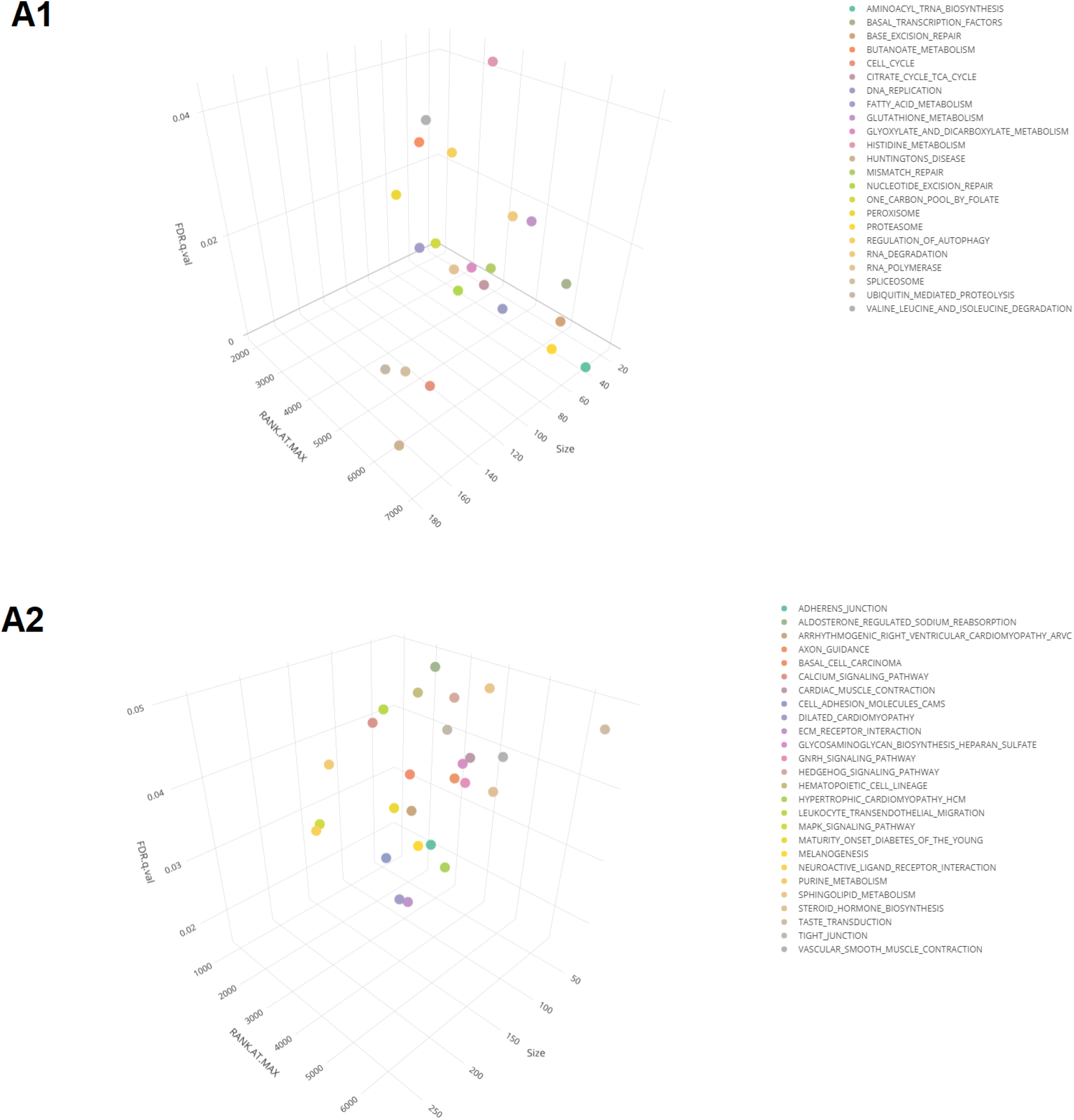
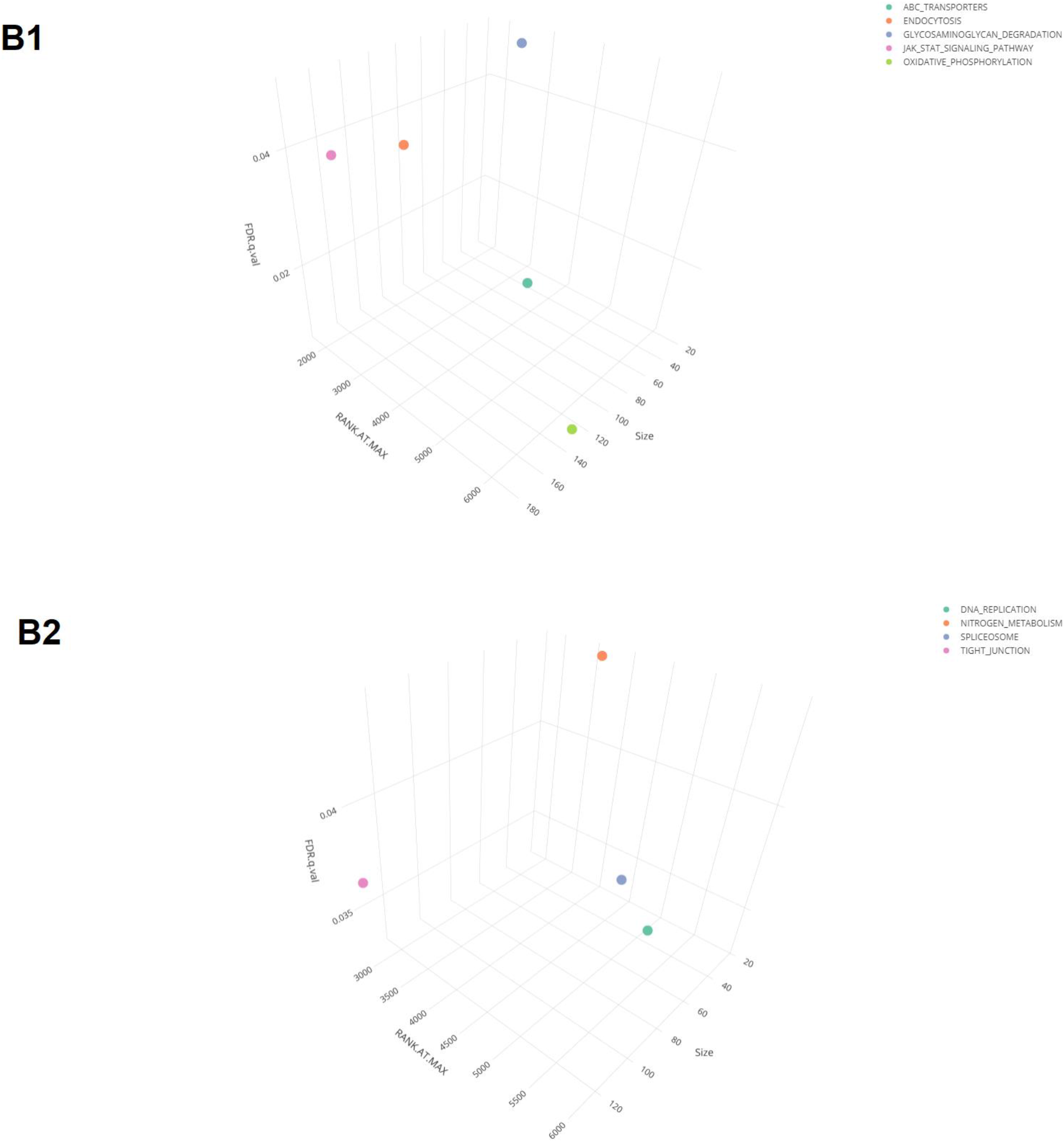
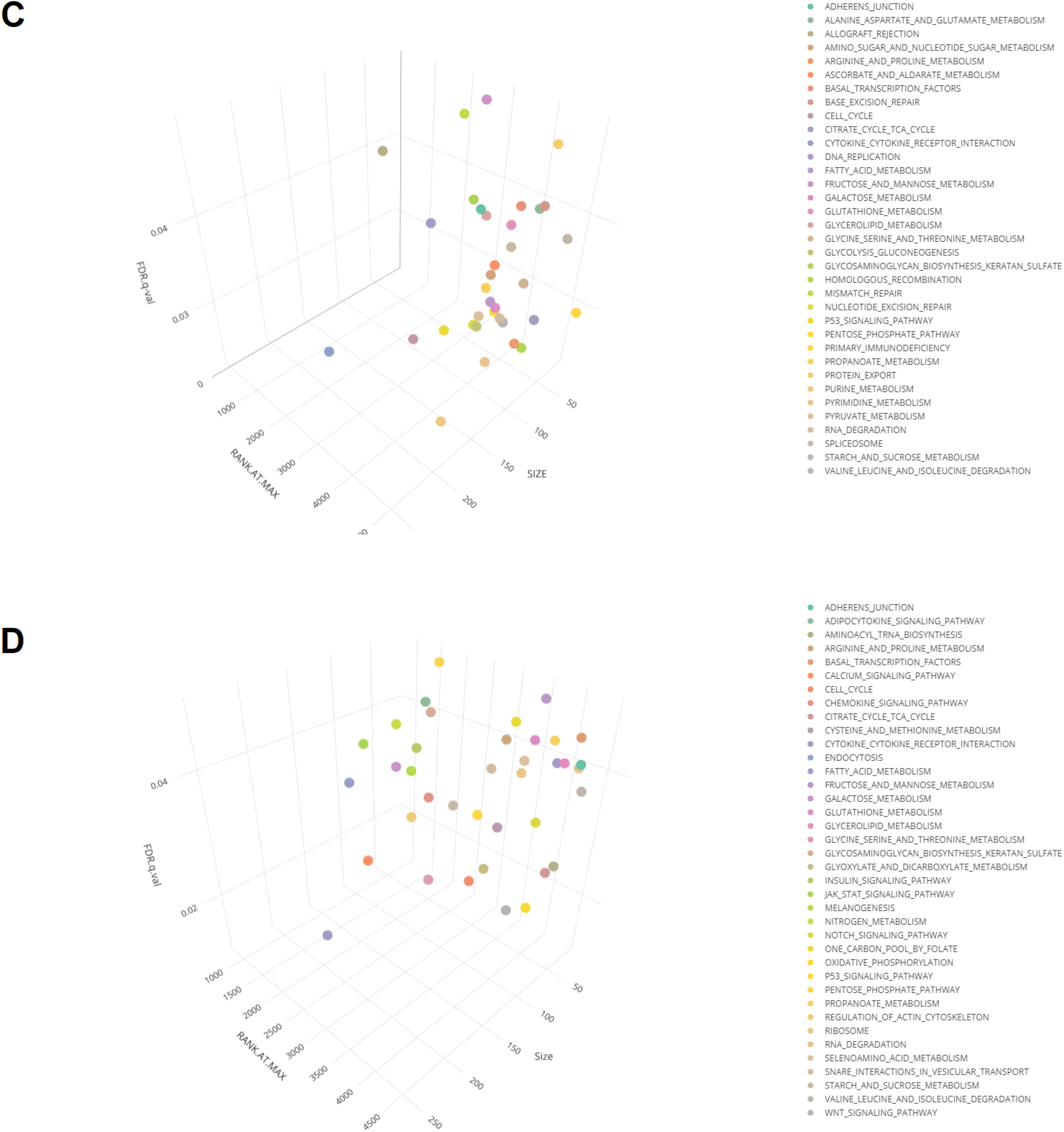
Cellular pathways.

**Figure 15.**
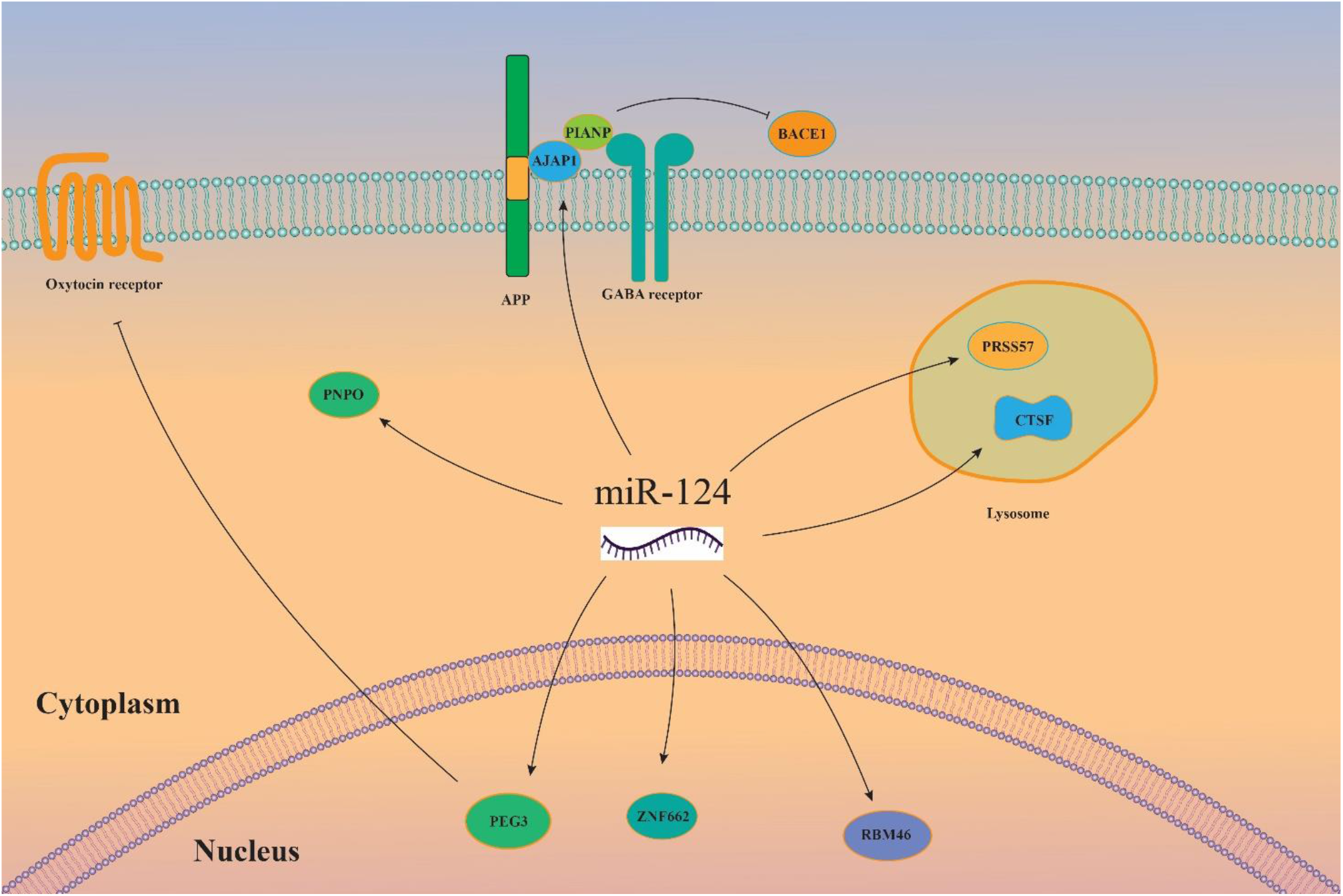
Overview of miR-124 function according RNA-seq data.

Many TFs expression is alliterated by inhibition of miR-124 and it is increased more over time (Figure 9). Therefore, we examined which TF could be targeted by miR-124. For this purpose, TargetScan was used and seven genes were fined, including NFIB, CDX2, DLX2, ELK3, NFIX, NR4A1, TAL1 find could target by miR-124 (Figure 10). NFIB is expressed in the neuronal progenitor cells line, which promote the specification of astrocytic and oligodendrocytic lineages (84). CDX2 is associated with NF-kB and inhibition of NF-kB activation decreases CDX2. Indeed, CDX2 has a role as downstream of NK-kB (85). DLX2 is associated with neuronal differentiation. Overexpression of DLX2 and ASCL1 induce neuronal differentiation, as well as GABAergic gene expression and electrophysiological maturity (86). ELK3 plays a role in the neuronal damage. ELK3 upregulates after peripheral neuronal damage and it seems that ELK3 can affect neuronal survival (87). NFIX contributes to neuronal proliferation and migration in the subventricular zone. NFIX^-/-^ mice have a higher number of neural progenitor cells in the subventricular zone but olfactory bulbs of NFIX^-/-^ mice are smaller (88). NR4A1 contributes to Aβ generation in the AD. NR4A1 increases in the hippocampus of APP/PS1 transgenic mice. Overexpression of NR4A1 in HT22 cells increased the levels of APP and BACE1. NR4A1 promotes amyloidogenic processing of APP in HT22 cells by regulating ADAM10 and BACE1 expression. Additionally, NR4A1 promote tau hyperphosphorylation by GSK3 signaling (89). TAL1 is an apoptosis factor and it plays its role by regulating of PTEN and eventually inhibiting PI3K signaling pathway (90).

miR-124 suppresses several cellular pathways, such as MAPK signaling pathway (Figure 14). Overexpression of miR-124 inhibits MAPK14 activation, and downregulates inflammatory cytokines expression (91). MAPK is a key target of chronic inflammatory diseases such as AD and MS. Activation of MAPK signaling in AD contributes to Tau phosphorylation, neurotoxicity neuroinflammation and synaptic dysfunction. MAPK overactivity also induces neurodegeneration in MS (92, 93). Transduction of miR-124 mimic significantly decreases proinflammatory cytokine level (94). Studies have shown that many cytokines caused inflammation in the AD, PD and ALS and cytokine-mediated inflammation significantly elevates in the cerebrospinal fluid of AD, PD and ALS. There is a strong link between some cytokines and ND (95).

In conclusion, miR-124 has a dynamic role in the CNS and regulates many gene and TFs. The most data and studies are about AD and PD, so the function of miR-124 on MS, HD and ALS needs further studies. miR-124 improves neuron survival in the ND and it could inhibit the oxidative stress and inflammation and apoptosis. Therefore, miR-124 can be a promising therapeutic approach for ND.

## Reference

1. Chen WW, Zhang X, Huang WJ. Role of neuroinflammation in neurodegenerative diseases (Review). Mol Med Rep. 2016;13(4):3391–6.

2. Gitler AD, Dhillon P, Shorter J. Neurodegenerative disease: models, mechanisms, and a new hope. Dis Model Mech. 2017;10(5):499–502.

3. Mantzavinos V, Alexiou A. Biomarkers for Alzheimer’s Disease Diagnosis. Curr Alzheimer Res. 2017;14(11):1149–54.

4. Mendiola-Precoma J, Berumen LC, Padilla K, Garcia-Alcocer G. Therapies for Prevention and Treatment of Alzheimer’s Disease. Biomed Res Int. 2016;2016:2589276.

5. Oladele JO, Oyeleke OM, Oladele OT, Olaniyan M. Neuroprotective mechanism of Vernonia amygdalina in a rat model of neurodegenerative diseases. Toxicol Rep. 2020;7:1223–32.

6. Cummings J, Feldman HH, Scheltens P. The “rights” of precision drug development for Alzheimer’s disease. Alzheimers Res Ther. 2019;11(1):76.

7. Tysnes OB, Storstein A. Epidemiology of Parkinson’s disease. J Neural Transm (Vienna). 2017;124(8):901–5.

8. Maher P. The Potential of Flavonoids for the Treatment of Neurodegenerative Diseases. Int J Mol Sci. 2019;20(12).

9. Ridolfi B, Abdel-Haq H. Neurodegenerative Disorders Treatment: The MicroRNA Role. Curr Gene Ther. 2017;17(5):327–63.

10. van den Berg MMJ, Krauskopf J, Ramaekers JG, Kleinjans JCS, Prickaerts J, Briedé JJ. Circulating microRNAs as potential biomarkers for psychiatric and neurodegenerative disorders. Prog Neurobiol. 2020;185:101732.

11. Angelopoulou E, Paudel YN, Piperi C. miR-124 and Parkinson’s disease: A biomarker with therapeutic potential. Pharmacol Res. 2019;150:104515.

12. Ludwig N, Leidinger P, Becker K, Backes C, Fehlmann T, Pallasch C, et al. Distribution of miRNA expression across human tissues. Nucleic Acids Res. 2016;44(8):3865–77.

13. Ghafouri-Fard S, Shoorei H, Bahroudi Z, Abak A, Majidpoor J, Taheri M. An update on the role of miR-124 in the pathogenesis of human disorders. Biomed Pharmacother. 2021;135:111198.

14. Jiang D, Gong F, Ge X, Lv C, Huang C, Feng S, et al. Neuron-derived exosomes-transmitted miR-124-3p protect traumatically injured spinal cord by suppressing the activation of neurotoxic microglia and astrocytes. J Nanobiotechnology. 2020;18(1):105.

15. Edgar R, Domrachev M, Lash AE. Gene Expression Omnibus: NCBI gene expression and hybridization array data repository. Nucleic Acids Res. 2002;30(1):207–10.

16. Kutsche LK, Gysi DM, Fallmann J, Lenk K, Petri R, Swiersy A, et al. Combined Experimental and System-Level Analyses Reveal the Complex Regulatory Network of miR-124 during Human Neurogenesis. Cell Syst. 2018;7(4):438–52.e8.

17. Lambert SA, Jolma A, Campitelli LF, Das PK, Yin Y, Albu M, et al. The Human Transcription Factors. Cell. 2018;172(4):650–65.

18. Subramanian A, Tamayo P, Mootha VK, Mukherjee S, Ebert BL, Gillette MA, et al. Gene set enrichment analysis: a knowledge-based approach for interpreting genome-wide expression profiles. Proc Natl Acad Sci U S A. 2005;102(43):15545–50.

19. Liberzon A, Subramanian A, Pinchback R, Thorvaldsdóttir H, Tamayo P, Mesirov JP. Molecular signatures database (MSigDB) 3.0. Bioinformatics. 2011;27(12):1739–40.

20. Xie Z, Bailey A, Kuleshov MV, Clarke DJB, Evangelista JE, Jenkins SL, et al. Gene Set Knowledge Discovery with Enrichr. Curr Protoc. 2021;1(3):e90.

21. Piñero J, Ramírez-Anguita JM, Saüch-Pitarch J, Ronzano F, Centeno E, Sanz F, et al. The DisGeNET knowledge platform for disease genomics: 2019 update. Nucleic Acids Res. 2020;48(D1):D845–d55.

22. Warde-Farley D, Donaldson SL, Comes O, Zuberi K, Badrawi R, Chao P, et al. The GeneMANIA prediction server: biological network integration for gene prioritization and predicting gene function. Nucleic Acids Res. 2010;38(Web Server issue):W214–20.

23. Li AD, Tong L, Xu N, Ye Y, Nie PY, Wang ZY, et al. miR-124 regulates cerebromicrovascular function in APP/PS1 transgenic mice via C1ql3. Brain Res Bull. 2019;153:214–22.

24. Hafez HA, Kamel MA, Osman MY, Osman HM, Elblehi SS, Mahmoud SA. Ameliorative effects of astaxanthin on brain tissues of alzheimer’s disease-like model: cross talk between neuronal-specific microRNA-124 and related pathways. Mol Cell Biochem. 2021;476(5):2233–49.

25. Yue D, Guanqun G, Jingxin L, Sen S, Shuang L, Yan S, et al. Silencing of long noncoding RNA XIST attenuated Alzheimer’s disease-related BACE1 alteration through miR-124. Cell Biol Int. 2020;44(2):630–6.

26. Zhao MY, Wang GQ, Wang NN, Yu QY, Liu RL, Shi WQ. The long-non-coding RNA NEAT1 is a novel target for Alzheimer’s disease progression via miR-124/BACE1 axis. Neurol Res. 2019;41(6):489–97.

27. Zhou Y, Deng J, Chu X, Zhao Y, Guo Y. Role of Post-Transcriptional Control of Calpain by miR-124-3p in the Development of Alzheimer’s Disease. J Alzheimers Dis. 2019;67(2):571–81.

28. Fernandes A, Ribeiro AR, Monteiro M, Garcia G, Vaz AR, Brites D. Secretome from SH-SY5Y APP(Swe) cells trigger time-dependent CHME3 microglia activation phenotypes, ultimately leading to miR-21 exosome shuttling. Biochimie. 2018;155:67–82.

29. Dalli T, Beker M, Terzioglu-Usak S, Akbas F, Elibol B. Thymoquinone activates MAPK pathway in hippocampus of streptozotocin-treated rat model. Biomed Pharmacother. 2018;99:391–401.

30. Zhang N, Zhao L, Su Y, Liu X, Zhang F, Gao Y. Syringin Prevents Aβ(25-35)-Induced Neurotoxicity in SK-N-SH and SK-N-BE Cells by Modulating miR-124-3p/BID Pathway. Neurochem Res. 2021;46(3):675–85.

31. Du X, Huo X, Yang Y, Hu Z, Botchway BOA, Jiang Y, et al. miR-124 downregulates BACE 1 and alters autophagy in APP/PS1 transgenic mice. Toxicol Lett. 2017;280:195–205.

32. Hou TY, Zhou Y, Zhu LS, Wang X, Pang P, Wang DQ, et al. Correcting abnormalities in miR-124/PTPN1 signaling rescues tau pathology in Alzheimer’s disease. J Neurochem. 2020;154(4):441–57.

33. Kang Q, Xiang Y, Li D, Liang J, Zhang X, Zhou F, et al. MiR-124-3p attenuates hyperphosphorylation of Tau protein-induced apoptosis via caveolin-1-PI3K/Akt/GSK3β pathway in N2a/APP695swe cells. Oncotarget. 2017;8(15):24314–26.

34. Feng CZ, Yin JB, Yang JJ, Cao L. Regulatory factor X1 depresses ApoE-dependent Aβ uptake by miRNA-124 in microglial response to oxidative stress. Neuroscience. 2017;344:217–28.

35. Kong Y, Wu J, Zhang D, Wan C, Yuan L. The Role of miR-124 in Drosophila Alzheimer’s Disease Model by Targeting Delta in Notch Signaling Pathway. Curr Mol Med. 2015;15(10):980–9.

36. Zhao YN, Li WF, Li F, Zhang Z, Dai YD, Xu AL, et al. Resveratrol improves learning and memory in normally aged mice through microRNA-CREB pathway. Biochem Biophys Res Commun. 2013;435(4):597–602.

37. Wang X, Wang ZH, Wu YY, Tang H, Tan L, Wang X, et al. Melatonin attenuates scopolamine-induced memory/synaptic disorder by rescuing EPACs/miR-124/Egr1 pathway. Mol Neurobiol. 2013;47(1):373–81.

38. Smith P, Al Hashimi A, Girard J, Delay C, Hébert SS. In vivo regulation of amyloid precursor protein neuronal splicing by microRNAs. J Neurochem. 2011;116(2):240–7.

39. Cao H, Han X, Jia Y, Zhang B. Inhibition of long non-coding RNA HOXA11-AS against neuroinflammation in Parkinson’s disease model via targeting miR-124-3p mediated FSTL1/NF-κB axis. Aging (Albany NY). 2021;13(8):11455–69.

40. Liu J, Liu D, Zhao B, Jia C, Lv Y, Liao J, et al. Long non-coding RNA NEAT1 mediates MPTP/MPP(+)-induced apoptosis via regulating the miR-124/KLF4 axis in Parkinson’s disease. Open Life Sci. 2020;15(1):665–76.

41. Ravanidis S, Bougea A, Papagiannakis N, Koros C, Simitsi AM, Pachi I, et al. Validation of differentially expressed brain-enriched microRNAs in the plasma of PD patients. Ann Clin Transl Neurol. 2020;7(9):1594–607.

42. Xing RX, Li LG, Liu XW, Tian BX, Cheng Y. Down regulation of miR-218, miR-124, and miR-144 relates to Parkinson’s disease via activating NF-κB signaling. Kaohsiung J Med Sci. 2020;36(10):786–92.

43. Lu Y, Gong Z, Jin X, Zhao P, Zhang Y, Wang Z. LncRNA MALAT1 targeting miR-124-3p regulates DAPK1 expression contributes to cell apoptosis in Parkinson’s Disease. J Cell Biochem. 2020.

44. Liu W, Zhang Q, Zhang J, Pan W, Zhao J, Xu Y. Long non-coding RNA MALAT1 contributes to cell apoptosis by sponging miR-124 in Parkinson disease. Cell Biosci. 2017;7:19.

45. Wang J, Wang W, Zhai H. MicroRNA-124 Enhances Dopamine Receptor Expression and Neuronal Proliferation in Mouse Models of Parkinson’s Disease via the Hedgehog Signaling Pathway by Targeting EDN2. Neuroimmunomodulation. 2019;26(4):174–87.

46. Gan L, Li Z, Lv Q, Huang W. Rabies virus glycoprotein (RVG29)-linked microRNA-124-loaded polymeric nanoparticles inhibit neuroinflammation in a Parkinson’s disease model. Int J Pharm. 2019;567:118449.

47. Geng L, Liu W, Chen Y. miR-124-3p attenuates MPP(+)-induced neuronal injury by targeting STAT3 in SH-SY5Y cells. Exp Biol Med (Maywood). 2017;242(18):1757–64.

48. Dong RF, Zhang B, Tai LW, Liu HM, Shi FK, Liu NN. The Neuroprotective Role of MiR-124-3p in a 6-Hydroxydopamine-Induced Cell Model of Parkinson’s Disease via the Regulation of ANAX5. J Cell Biochem. 2018;119(1):269–77.

49. Saraiva C, Ferreira L, Bernardino L. Traceable microRNA-124 loaded nanoparticles as a new promising therapeutic tool for Parkinson’s disease. Neurogenesis (Austin). 2016;3(1):e1256855.

50. Li N, Pan X, Zhang J, Ma A, Yang S, Ma J, et al. Plasma levels of miR-137 and miR-124 are associated with Parkinson’s disease but not with Parkinson’s disease with depression. Neurol Sci. 2017;38(5):761–7.

51. Gong X, Wang H, Ye Y, Shu Y, Deng Y, He X, et al. miR-124 regulates cell apoptosis and autophagy in dopaminergic neurons and protects them by regulating AMPK/mTOR pathway in Parkinson’s disease. Am J Transl Res. 2016;8(5):2127–37.

52. Saraiva C, Paiva J, Santos T, Ferreira L, Bernardino L. MicroRNA-124 loaded nanoparticles enhance brain repair in Parkinson’s disease. J Control Release. 2016;235:291–305.

53. Wang H, Ye Y, Zhu Z, Mo L, Lin C, Wang Q, et al. MiR-124 Regulates Apoptosis and Autophagy Process in MPTP Model of Parkinson’s Disease by Targeting to Bim. Brain Pathol. 2016;26(2):167–76.

54. Zha Z, Gao YF, Ji J, Sun YQ, Li JL, Qi F, et al. Bu Shen Yi Sui Capsule Alleviates Neuroinflammation and Demyelination by Promoting Microglia toward M2 Polarization, Which Correlates with Changes in miR-124 and miR-155 in Experimental Autoimmune Encephalomyelitis. Oxid Med Cell Longev. 2021;2021:5521503.

55. Amoruso A, Blonda M, Gironi M, Grasso R, Di Francescantonio V, Scaroni F, et al. Immune and central nervous system-related miRNAs expression profiling in monocytes of multiple sclerosis patients. Sci Rep. 2020;10(1):6125.

56. Gandy KAO, Zhang J, Nagarkatti P, Nagarkatti M. Resveratrol (3, 5, 4’-Trihydroxy-trans-Stilbene) Attenuates a Mouse Model of Multiple Sclerosis by Altering the miR-124/Sphingosine Kinase 1 Axis in Encephalitogenic T Cells in the Brain. J Neuroimmune Pharmacol. 2019;14(3):462–77.

57. Malhotra S, Villar LM, Costa C, Midaglia L, Cubedo M, Medina S, et al. Circulating EZH2-positive T cells are decreased in multiple sclerosis patients. J Neuroinflammation. 2018;15(1):296.

58. Dutta R, Chomyk AM, Chang A, Ribaudo MV, Deckard SA, Doud MK, et al. Hippocampal demyelination and memory dysfunction are associated with increased levels of the neuronal microRNA miR-124 and reduced AMPA receptors. Ann Neurol. 2013;73(5):637–45.

59. Yelick J, Men Y, Jin S, Seo S, Espejo-Porras F, Yang Y. Elevated exosomal secretion of miR-124-3p from spinal neurons positively associates with disease severity in ALS. Exp Neurol. 2020;333:113414.

60. Lee ST, Im W, Ban JJ, Lee M, Jung KH, Lee SK, et al. Exosome-Based Delivery of miR-124 in a Huntington’s Disease Model. J Mov Disord. 2017;10(1):45–52.

61. Das E, Jana NR, Bhattacharyya NP. MicroRNA-124 targets CCNA2 and regulates cell cycle in STHdh(Q111)/Hdh(Q111) cells. Biochem Biophys Res Commun. 2013;437(2):217–24.

62. Jorda A, Cauli O, Santonja JM, Aldasoro M, Aldasoro C, Obrador E, et al. Changes in Chemokines and Chemokine Receptors Expression in a Mouse Model of Alzheimer’s Disease. Int J Biol Sci. 2019;15(2):453–63.

63. Kang Y, Xie H, Zhao C. Ankrd45 Is a Novel Ankyrin Repeat Protein Required for Cell Proliferation. Genes (Basel). 2019;10(6).

64. Broad KD, Curley JP, Keverne EB. Increased apoptosis during neonatal brain development underlies the adult behavioral deficits seen in mice lacking a functional paternally expressed gene 3 (Peg3). Dev Neurobiol. 2009;69(5):314–25.

65. Zhang M, Zhou X, Jiang W, Li M, Zhou R, Zhou S. AJAP1 affects behavioral changes and GABA(B)R1 level in epileptic mice. Biochem Biophys Res Commun. 2020;524(4):1057–63.

66. Gao C, Fu Q, Su B, Song H, Zhou S, Tan F, et al. The involvement of cathepsin F gene (CTSF) in turbot (Scophthalmus maximus L.) mucosal immunity. Fish Shellfish Immunol. 2017;66:270–9.

67. Bras J, Djaldetti R, Alves AM, Mead S, Darwent L, Lleo A, et al. Exome sequencing in a consanguineous family clinically diagnosed with early-onset Alzheimer’s disease identifies a homozygous CTSF mutation. Neurobiol Aging. 2016;46:236.e1–6.

68. Song Q, Zhao F, Yao J, Dai H, Hu L, Yu S. Protective effect of microRNA-134-3p on multiple sclerosis through inhibiting PRSS57 and promotion of CD34(+) cell proliferation in rats. J Cell Biochem. 2020;121(11):4347–63.

69. Zhai L, Wang C, Chen Y, Zhou S, Li L. Rbm46 regulates mouse embryonic stem cell differentiation by targeting β-Catenin mRNA for degradation. PLoS One. 2017;12(2):e0172420.

70. Kritsilis M, S VR, Koutsoudaki PN, Evangelou K, Gorgoulis VG, Papadopoulos D. Ageing, Cellular Senescence and Neurodegenerative Disease. Int J Mol Sci. 2018;19(10).

71. Fang P, Kazmi SA, Jameson KG, Hsiao EY. The Microbiome as a Modifier of Neurodegenerative Disease Risk. Cell Host Microbe. 2020;28(2):201–22.

72. Armstrong R. What causes neurodegenerative disease? Folia Neuropathol. 2020;58(2):93–112.

73. Juźwik CA, S SD, Zhang Y, Paradis-Isler N, Sylvester A, Amar-Zifkin A, et al. microRNA dysregulation in neurodegenerative diseases: A systematic review. Prog Neurobiol. 2019;182:101664.

74. Hampel H, Vassar R, De Strooper B, Hardy J, Willem M, Singh N, et al. The β-Secretase BACE1 in Alzheimer’s Disease. Biol Psychiatry. 2021;89(8):745–56.

75. Uruno A, Matsumaru D, Ryoke R, Saito R, Kadoguchi S, Saigusa D, et al. Nrf2 Suppresses Oxidative Stress and Inflammation in App Knock-In Alzheimer’s Disease Model Mice. Mol Cell Biol. 2020;40(6).

76. Gao Y, Tan L, Yu JT, Tan L. Tau in Alzheimer’s Disease: Mechanisms and Therapeutic Strategies. Curr Alzheimer Res. 2018;15(3):283–300.

77. Chen J, Wang X, Yi X, Wang Y, Liu Q, Ge R. Induction of KLF4 contributes to the neurotoxicity of MPP + in M17 cells: a new implication in Parkinson’s disease. J Mol Neurosci. 2013;51(1):109–17.

78. Pajares M, A IR, Manda G, Boscá L, Cuadrado A. Inflammation in Parkinson’s Disease: Mechanisms and Therapeutic Implications. Cells. 2020;9(7).

79. Frey WD, Sharma K, Cain TL, Nishimori K, Teruyama R, Kim J. Oxytocin receptor is regulated by Peg3. PLoS One. 2018;13(8):e0202476.

80. Mazurek MF, Beal MF, Bird ED, Martin JB. Oxytocin in Alzheimer’s disease: postmortem brain levels. Neurology. 1987;37(6):1001–3.

81. Dinamarca MC, Raveh A, Schneider A, Fritzius T, Früh S, Rem PD, et al. Complex formation of APP with GABA(B) receptors links axonal trafficking to amyloidogenic processing. Nat Commun. 2019;10(1):1331.

82. van der Zee J, Mariёn P, Crols R, Van Mossevelde S, Dillen L, Perrone F, et al. Mutated CTSF in adult-onset neuronal ceroid lipofuscinosis and FTD. Neurol Genet. 2016;2(5):e102.

83. Ciapaite J, Albersen M, Savelberg SMC, Bosma M, Tessadori F, Gerrits J, et al. Pyridox(am)ine 5’-phosphate oxidase (PNPO) deficiency in zebrafish results in fatal seizures and metabolic aberrations. Biochim Biophys Acta Mol Basis Dis. 2020;1866(3):165607.

84. Matuzelski E, Harvey TJ, Harkins D, Nguyen T, Ruitenberg MJ, Piper M. Expression of NFIA and NFIB within the murine spinal cord. Gene Expr Patterns. 2020;35:119098.

85. Yu JH, Zheng JB, Qi J, Yang K, Wu YH, Wang K, et al. Bile acids promote gastric intestinal metaplasia by upregulating CDX2 and MUC2 expression via the FXR/NF-κB signalling pathway. Int J Oncol. 2019;54(3):879–92.

86. Barretto N, Zhang H, Powell SK, Fernando MB, Zhang S, Flaherty EK, et al. ASCL1- and DLX2-induced GABAergic neurons from hiPSC-derived NPCs. J Neurosci Methods. 2020;334:108548.

87. Kerr N, Pintzas A, Holmes F, Hobson SA, Pope R, Wallace M, et al. The expression of ELK transcription factors in adult DRG: Novel isoforms, antisense transcripts and upregulation by nerve damage. Mol Cell Neurosci. 2010;44(2):165–77.

88. Heng YH, Zhou B, Harris L, Harvey T, Smith A, Horne E, et al. NFIX Regulates Proliferation and Migration Within the Murine SVZ Neurogenic Niche. Cereb Cortex. 2015;25(10):3758–78.

89. Zhao LG, Tang Y, Tan JZ, Wang JW, Chen GJ, Zhu BL. The effect of NR4A1 on APP metabolism and tau phosphorylation. Genes Dis. 2018;5(4):342–8.

90. Wu Y, Hu Y, Yu X, Zhang Y, Huang X, Chen S, et al. TAL1 mediates imatinib-induced CML cell apoptosis via the PTEN/PI3K/AKT pathway. Biochem Biophys Res Commun. 2019;519(2):234–9.

91. Pan W, Wei N, Xu W, Wang G, Gong F, Li N. MicroRNA-124 alleviates the lung injury in mice with septic shock through inhibiting the activation of the MAPK signaling pathway by downregulating MAPK14. Int Immunopharmacol. 2019;76:105835.

92. Lee JK, Kim NJ. Recent Advances in the Inhibition of p38 MAPK as a Potential Strategy for the Treatment of Alzheimer’s Disease. Molecules. 2017;22(8).

93. Ten Bosch GJA, Bolk J, t Hart BA, Laman JD. Multiple sclerosis is linked to MAPK(ERK) overactivity in microglia. J Mol Med (Berl). 2021;99(8):1033–42.

94. Yu A, Zhang T, Duan H, Pan Y, Zhang X, Yang G, et al. MiR-124 contributes to M2 polarization of microglia and confers brain inflammatory protection via the C/EBP-α pathway in intracerebral hemorrhage. Immunol Lett. 2017;182:1–11.

95. Chen X, Hu Y, Cao Z, Liu Q, Cheng Y. Cerebrospinal Fluid Inflammatory Cytokine Aberrations in Alzheimer’s Disease, Parkinson’s Disease and Amyotrophic Lateral Sclerosis: A Systematic Review and Meta-Analysis. Front Immunol. 2018;9:2122.

